# Unpacking the functional heterogeneity of the Emotional Face Matching Task: a normative modelling approach

**DOI:** 10.1101/2023.03.27.534351

**Authors:** Hannah S. Savage, Peter C. R. Mulders, Philip F. P. van Eijndhoven, Jasper van Oort, Indira Tendolkar, Janna N. Vrijsen, Christian F. Beckmann, Andre F. Marquand

**Affiliations:** Donders Institute of Brain, Cognition and Behaviour, Radboud University, Nijmegen, The Netherlands; Department of Cognitive Neuroscience, Radboud University Medical Centre, Nijmegen, The Netherlands; Department of Psychiatry, Radboud University Medical Centre, Nijmegen, The Netherlands; Depression Expertise Centre, Pro Persona Mental Health Care, Nijmegen, The Netherlands; Centre for Functional MRI of the Brain (FMRIB), Nuffield Department of Clinical Neurosciences, Wellcome Centre for Integrative Neuroimaging, University of Oxford, Oxford, UK

## Abstract

Functional neuroimaging has contributed substantially to understanding brain function but is dominated by group analyses that index only a fraction of the variation in these data. It is increasingly clear that parsing the underlying heterogeneity is crucial to understand individual differences and the impact of different task manipulations. We estimate large-scale (N=7728) normative models of task-evoked activation during the Emotional Face Matching Task, which enables us to bind heterogeneous datasets to a common reference and dissect heterogeneity underlying group-level analyses. We apply this model to a heterogenous patient cohort, to map individual differences between patients with one or more mental health diagnoses relative to the reference cohort and determine multivariate associations with transdiagnostic symptom domains. For the face>shapes contrast, patients have a higher frequency of extreme deviations which are spatially heterogeneous. In contrast, normative models for faces>baseline have greater predictive value for individuals’ transdiagnostic functioning.

## Introduction

Task-based functional neuroimaging (functional magnetic resonance imaging; fMRI) has been widely applied in foundational and clinical neuropsychology to characterise neural processes that underpin a behaviour or process of interest. The typical approach in such studies is based on comparing mean differences in the magnitude and location of activation (measured by changes in BOLD signal), which has helped us to understand how these processes may differ between groups defined by biological and sociocultural factors, psychopathologies, or therapeutic interventions. The majority of prior research has reported group-level summary statistics, which inform us of those regions most consistently activated across participants/groups during task conditions. This method assumes that the neural mechanisms facilitating the process of interest are consistent across individuals within and between groups. This assumption enables our understanding to reach only so far as ‘the average brain’ of an ‘average control’, or ‘average patient’.

In order to better understand how the brain relates to behaviour it is essential to move our focus from the group-level to studying individual differences and consider the neural activation of these processes within the context of multiple sources of heterogeneity. For example: (i) natural variation within the general population, including potentially heterogenous yet functionally convergent processes, and (ii) heterogeneity within groups of interest, such as within mental health diagnoses. Furthermore, when comparing between independent studies, the influence of task design (i.e. small modifications to an original task) and acquisition parameters should also be considered but are seldom investigated.

One approach that can provide insight into individual differences is normative modelling^1,2^. The normative modelling framework provides statistical inference at the level of each subject with respect to an expected pattern across the population, highlighting variation within populations in terms of the mapping between biological variables and other measures of interest. This framework has previously been employed by our group and others to dissect structural variation within large healthy populations^3^ and clinical psychiatric populations (e.g. in autism ^4–6^, schizophrenia and bipolar disorder^7^), and in relation to dimensions of psychopathology^8^. Applying this method to task-based fMRI data we will be able to characterize how functional activity within each voxel or ROI in the brain differs between individuals, and hence show with greater nuance the range of task-evoked activation within the general population^2^. Further, applying this model to patients with a current diagnosis (mood and anxiety disorders, autism spectrum disorders (ASD) and/or attention deficit hyperactivity disorder (ADHD)) we will be able to map differences in these individual participants with respect to the reference cohort. This may reveal unique clusters of deviation patterns, within and/or across diagnostic categories.

In this study, we use the Emotional Face Matching Task (EFMT) to demonstrate the potential of the normative modelling method to identify individual differences in task-based fMRI. The EFMT, also commonly referred to as the ‘Hariri task’, has been used in over 250 fMRI studies since it was most notably introduced in 2002^9,10^. This task asks participants to match one of two images that are simultaneously presented at the bottom of the screen, to a third target image displayed at the top of the screen; participants match images of facial configurations consistent with the common view of prototypic facial expressions, most frequently of fear or anger, or similarly positioned geometric shapes. Matching faces, as compared to matching shapes, evokes explicit and/or implicit emotional face processing, which has previously been shown to engage the amygdala, fusiform face area, anterior insula cortex, the pregenual and dorsal anterior cingulate cortex, the dorsomedial and dorsolateral prefrontal cortex, and visual input areas. Previous work has related activity to biological and demographic variables, and compared between many different clinical groups and developmental spectrums.

Due to its experimental simplicity and focus on subcortical circuitry relevant to brain disorders, the EFMT has been implemented in a number of large-scale neuroimaging initiatives including the UK Biobank^11^, the Human Connectome Project (HCP)^12,13^, HCP Development^14^, the Amsterdam Open MRI Collection Population Imaging of Psychology (AOMIC PIOP2)^15^, and the Duke Neurogenetics Study (DNS). We take advantage of these large open-access/shared datasets to collate a large representative sample of over 7500 participants from six sites to first (1) build reference normative models that highlight the natural variation of functional activity evoked by the EFTM [as measured by the task contrasts faces > shapes and faces>baseline], and (2) determine how the model’s prediction relates to age, sex, and variations in task design. We then apply these models to over 200 participants with a current mental health condition or who are neurodivergent from the MIND-Set cohort (Measuring Integrated Novel Dimensions in neurodevelopmental and stress-related psychiatric disorder)^16^, to (3) map deviations in patients with a current diagnosis (mood and anxiety disorders, ASD and/or ADHD) relative from the reference cohort, both at the group level and at the level of the individual. We show that despite the ostensible simplicity of this task and robust group effects, there is considerable inter-individual heterogeneity in the nature of the elicited activation patterns and that such differences are both highly interpretable and predict cross-domain symptomatology in a naturalistic clinical cohort.

## Results

### Group level comparisons show consistent effects across cohorts

First, we performed a classical group comparison to provide a reference against which to understand the inter-individual differences in subsequent analyses. To achieve this, we randomly selected 100 random individuals’ FSL pre-processed data into fixed-effects general linear models to create group level maps for the faces>shapes (Fig. 1a) and faces>baseline (Fig. 1b) contrasts (see methods). This also served as a sanity check to ensure the data was comparable to past literature. Overall, positive task effects (activations) for faces>shapes were found in the bilateral inferior and middle occipital lobe and the calcarine cortex (V1) extending anterior-ventrally to the bilateral lingual and fusiform gyrus, and anterior-dorsally to the middle and inferior temporal gyrus; the bilateral amygdala extending into the hippocampus; the bilateral temporal pole; a dorsal region of the vmPFC; and the bilateral middle and inferior frontal gyrus. Task-related deactivations were found across regions comprising the default mode network, including the anterior and posterior cingulate cortex and precuneus, the precentral gyrus and supplementary motor area and the inferior temporal lobe.

**Figure 1:**
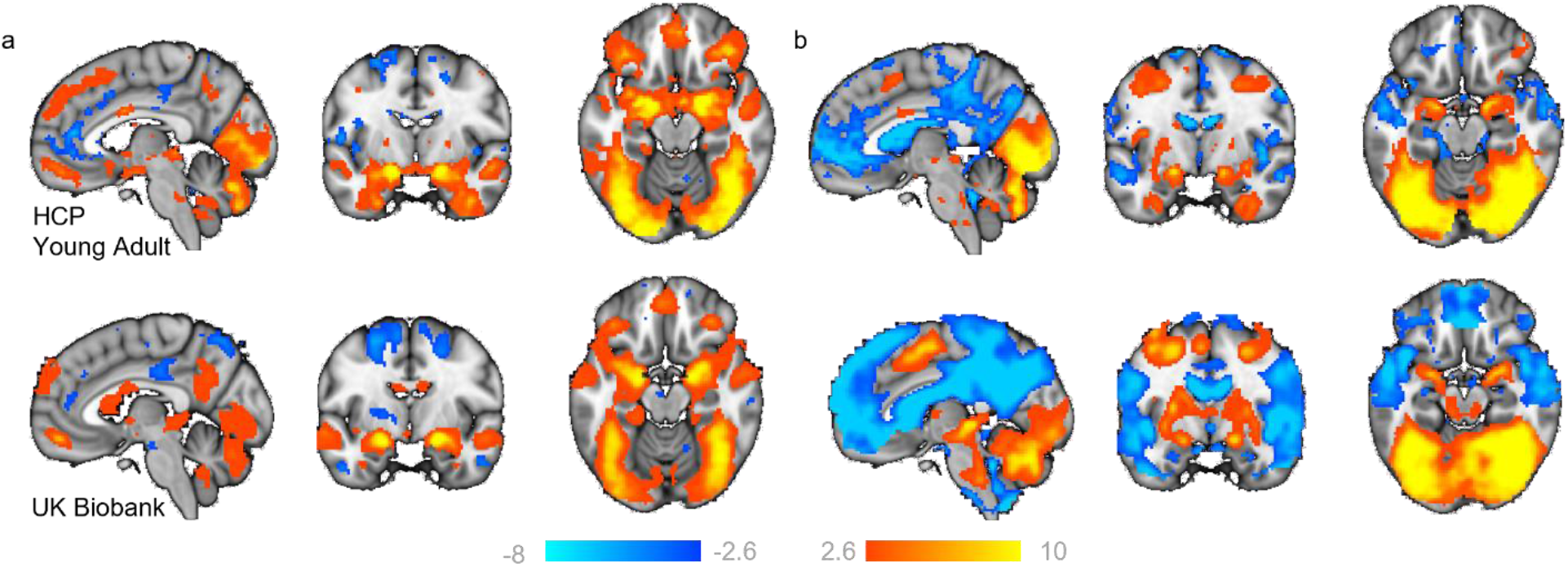
Task evoked activation. Two representative groups maps (from HCP Young Adult and UK Biobank), illustrating regions where participants show greater BOLD signal (z-statistic maps, thresholded at>±2.6) to (a) faces, as compared to shapes (faces>shapes), and (b) faces, as compared to baseline (faces>baseline). x,y,z = -4,-6,-16.

### Fitting reference normative models for emotional face processing

Next, we estimated normative models of EFMT-related BOLD activation for the face>shapes and faces>baseline contrast using data from 7728 individuals across the lifespan. To achieve this, we split the data into training and test splits, stratified by site (face>shapes – train: 3885, test: 3843; faces>baseline – train: 3778, test: 3950; see Supplementary Figure 1), then fit a Bayesian Linear Regression model that predicted the single subject level activation for each voxel of the brain, as a function of sex, age, and acquisition and task parameters (see methods). Explained variance in the test set was good (reaching 0.525), especially in regions that showed activation at the group level (Fig. 2) including the occipital lobe/visual cortex and the bilateral amygdala (faces>shapes: Fig. 2a; faces>baseline: Fig. 2d). As shown in Supplementary Fig. 2a and 2c in most voxels the skew and kurtosis was acceptable (i.e. -1 < skew < 1 and kurtosis around zero). For a very small proportion of voxels this was not the case; the most ventral region of the vmPFC (i.e. the bottom border of the brain) was the most negatively skewed. As these voxels spatially overlap with those showing positive kurtosis, which likely reflects the extended negative tails of the distributions in these voxels, we interpret this to reflect the varying degrees of signal dropout, more so than biological variation. Despite our efforts to ensure signal coverage within these voxels and minimal motion artefacts in the data used to construct the model, we do advise readers to interpret deviation scores within this region with caution.

**Figure 2:**
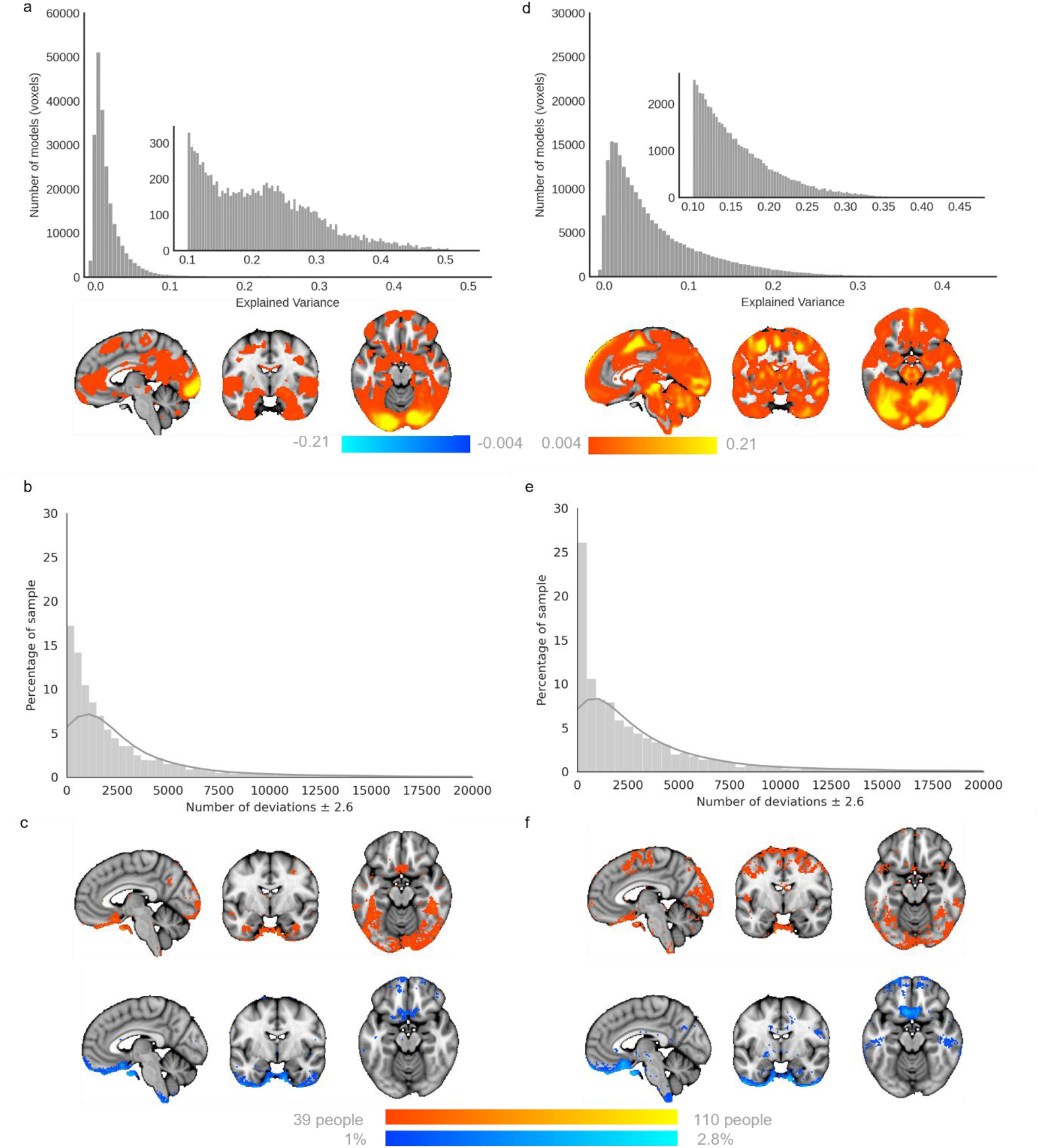
Evaluation and deviation scores from the faces>shapes (left) and faces>baseline (right) normative models. Explained variance is high in the normative models, irrespective of whether they are built using the face>shapes contrast (a), or the faces>baseline contrast. (d). Histograms show the relative frequency of the total number of deviations that a participant has for each model (b,e). Normative Probability Maps illustrate the percentage of participants of the total sample who had positive (hot colours) or negative deviations (cool colours) >±2.6 within each voxel, for the faces > shapes (c) and faces>baseline (f) models. x,y,z = -4,-6,-16

### Voxel-wise deviations show considerable inter-individual variability

We then used these normative models to quantify the degree of inter-individual variability. To achieve this, for each participant we created a thresholded normative probability map (NPM; deviation scores >±2.6) which indicates the difference between the predicted activation and true activation scaled by the prediction variance, and therefore shows the voxels where that participant had greater or less activation than would be expected by the normative models. Figures 2b and 2e show the frequency of the total number of deviations that individuals had from the faces>shapes, and faces>baseline models, respectively. Within each voxel, we then counted how many participants had positive or negative deviations (>±2.6). The resulting brain maps illustrate the variability in the magnitude of functional activation per voxel, across the population for the two task contrasts (Fig. 2c + f). This shows that: (i) there is considerable inter-individual variability underlying the mean effects and (ii) that the spatial distribution of individual deviations mostly occurs within the task network. Every voxel of the brain had at least one subject with a deviation >±2.6 (not shown), although, as illustrated, there were regions including the medial occipital lobe extending to the bilateral fusiform gyrus and inferior temporal lobe, the bilateral inferior frontal gyrus extending to the precentral gyrus, and the posterior region of the vmPFC, wherein deviations were more frequently observed. As there were minimal differences in the evaluation metrics between models built using either contrast, and as the contrast faces>shapes is most commonly reported in prior literature, we use this as our primary contrast for our further analysis of the reference model.

### Voxel-wise deviations are reliable: Test-Retest

The aforementioned normative models were re-generated removing all participants from the original HCP Young Adult sample for whom HCP Retest data was available (n = 42). The original HCP Young Adult (Test) data and the Retest data were then independently applied to generate voxel-wise deviation scores per individual, per session. The magnitude of deviations from Test and Retest sessions were highly correlated, especially in regions including the medial occipital lobe extending to the bilateral fusiform gyrus and inferior temporal lobe wherein large deviations were more frequently observed. Deviation scores in only 5.2% of voxels were significantly different between Test and Retest, with the largest differences (>±0.5) predominantly observed within the vmPFC region. This is consistent with the notion that deviations within this region are particularly sensitive to signal drop out which is likely session dependent; synonymously, deviations in this region were also not highly correlated. Qualitatively, the NPMs were replicated across Test-Retest sessions. This analysis was also performed for the faces>baseline models; results lead to the same conclusions as for the face>shapes models (Supplementary Figure 5).

### Separable effects of input variables on model predictions

Next, we examined structure coefficients from our models to disentangle the effects of different input variables. Structure coefficients provide insight into the bivariate relationship between the effect observed (in this case the predicted z-stat BOLD activation), and the predictor (or covariate of interest) without the influence of other covariates in the model. As shown in Figure 3, the direction of the relationship between input variables and the predicted BOLD activation, and the fraction of the explained variability can be meaningfully separated for interpretation. Some input variables, namely acquisition parameters, showed overlapping effects (with sensical direction flips) likely due to their relatively high correlation and limited variability across sites; the number of target blocks, volumes acquired (not shown), use of multiband sequence (not shown), the length of the TR (not shown), and site (not shown) all showed a similar relation to predicted activity.

**Figure 3.**
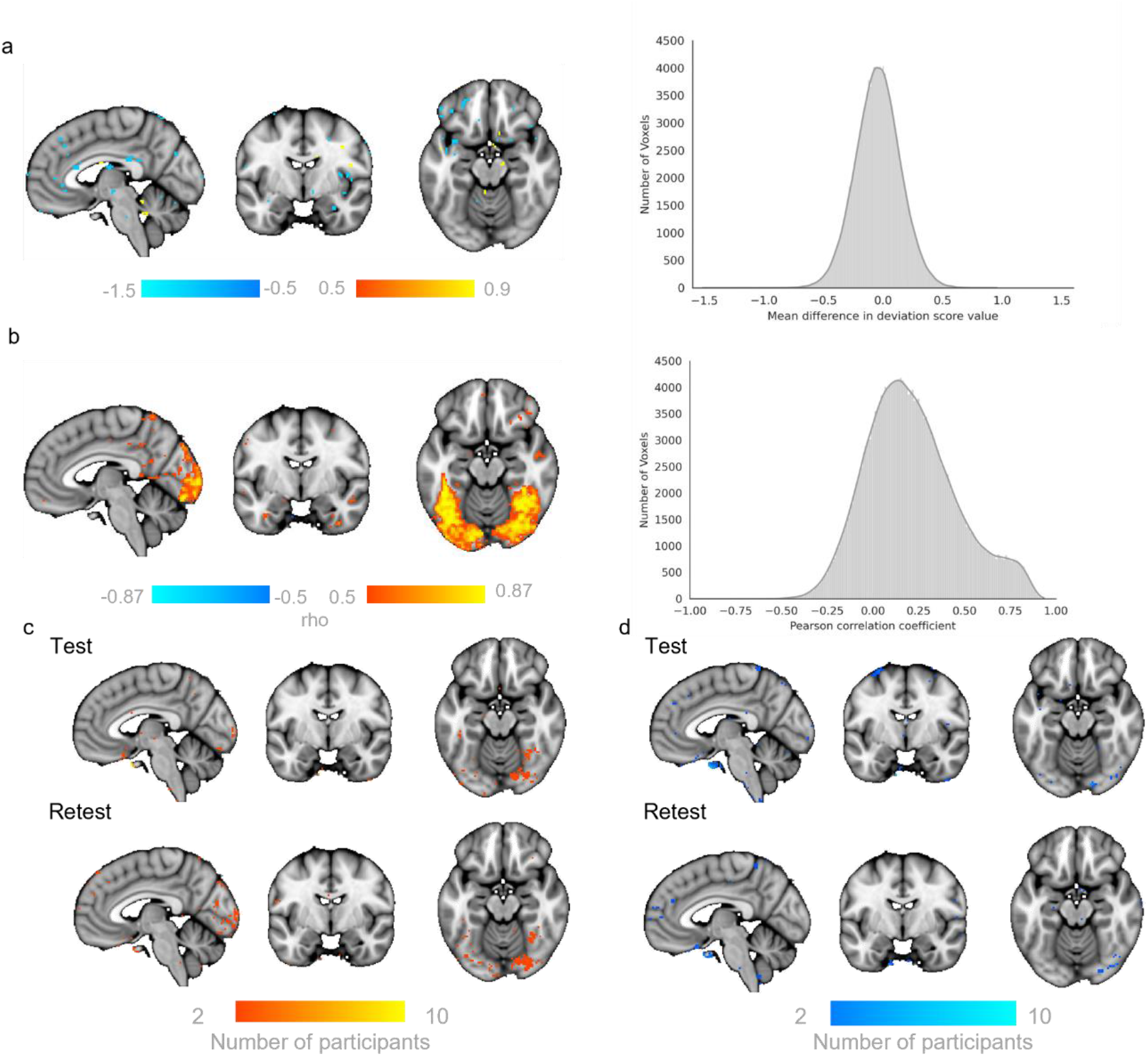
Test – Retest reliability of deviation scores for faces>shapes models. Mean within-subject difference per voxel (histogram) illustrated thresholded at >0.5 (i.e. a change greater than half a standard deviation between Test and Retest scans (a). The correlation coefficients (rho) between Test and Retest deviation scores (histogram) illustrated thresholded by the coefficients of determination (rho^2^>0.3, b). Normative Probability Maps illustrate the voxels wherein 1 or more participant had positive (c, hot colours) or negative deviations (d, cool colours) > ±2.6 for the faces > shapes normative models in the Test (top rows) and Re-Test (bottom rows) samples. x,y,z = -4,-6,-16

Increased age (Fig. 3. Top row - Age) was related to decreased predicted activity across the peripheral/surface of the brain, as well as regions surrounding the ventricles, and increased activity in midline regions of the default mode network, the bilateral insula, the fusiform face area extending to the para-hippocampal gyrus and the superior temporal gyrus. Being female (Fig. 3. Top row - Sex), was related to increased mid-to-anterior insula and cingulate cortex activation. Predictions were only minimally influenced by intra-cranial volume (not shown).

We further illustrate the ability of this method to disentangle the influence of task design choices, on predicted activation. For example, task length, the influence of the matching rule and the stimuli presented. The longer the task (Fig 3. Middle row - Task length), the greater the activation within the bilateral amygdala, bilateral insula and V2. Being told to match the emotional expression, as compared to matching the faces, related to increased predicted BOLD activity within subcortical areas including the bilateral putamen, caudate body and medio-dorsal thalamus (Fig. 3. Bottom row - Instructions). Attending to the emotional expression also predicted increased activity within the mid-cingulate and superior frontal gyrus extending to the supplementary motor area, the posterior medial temporal gyrus the inferior temporal gyrus, and the medial temporal pole. Conversely, when participants were asked to match faces (Fig. 3. Bottom row - Instructions), the model predicted greater activation within the bilateral fusiform gyrus, the middle temporal gyrus, the superior temporal pole, the dorsolateral prefrontal cortex, and a large area of the inferior parietal gyrus extending to the supramarginal and angular gyrus. Additionally, when stimuli from the Ekman series were used (Fig. 3. Middle row - Target stimuli) the model predicted greater activation within the bilateral inferior occipital gyrus and the calcarine cortex (V1), the bilateral lingual and fusiform gyrus extending to the inferior temporal gyrus, as well as in the medial cingulate cortex, an anterior region of the vmPFC, the superior medial prefrontal cortex, and subcortical regions including the ventral posterior thalamus, the posterior putamen, para-hippocampus, hippocampus and amygdala. Conversely, the use of the Nim-Stim Set stimuli related to greater activity within default mode regions, including a large area of the ventromedial/medial prefrontal cortex, precuneus, cuneus, as well as the supramarginal gyrus which extended medially to the anterior and posterior insula, which in turn extended laterally to the superior and medial temporal gyri (Fig. 3. Middle row - Target stimuli).

### A traditional case-control comparison identifies few differences between patients and controls

We then performed a voxel-wise case-control comparison on the raw data to test for group level differences between a heterogeneous patient cohort and matched unaffected controls from the naturalistic MIND-Set sample. As evidenced in Table 1 (see Diagnoses), the naturalistic MIND-Set sample has many patients with co-occurring and heterogenous mental health diagnosis, with or without neurodivergence, and is therefore representative of diverse clinical populations. This analysis revealed very few differences between the patient cohort, and unaffected controls for faces>shapes and faces>baseline (Fig. 4a and b – bottom rows). More specifically, comparing patients’ task activation (Fig. 4a – top row) to controls (Fig. 4a – middle row) for the faces>shapes contrast showed patients had greater activation in the left temporal medial gyrus and bilateral posterior cingulate cortex, as well as in small regions of the supplementary motor area, and the genus of the anterior cingulate cortex (Fig. 4a – bottom row). There were negligible differences between patients and unaffected controls for the faces>baseline contrast (Fig. 4b – bottom row).

**Figure 4:**
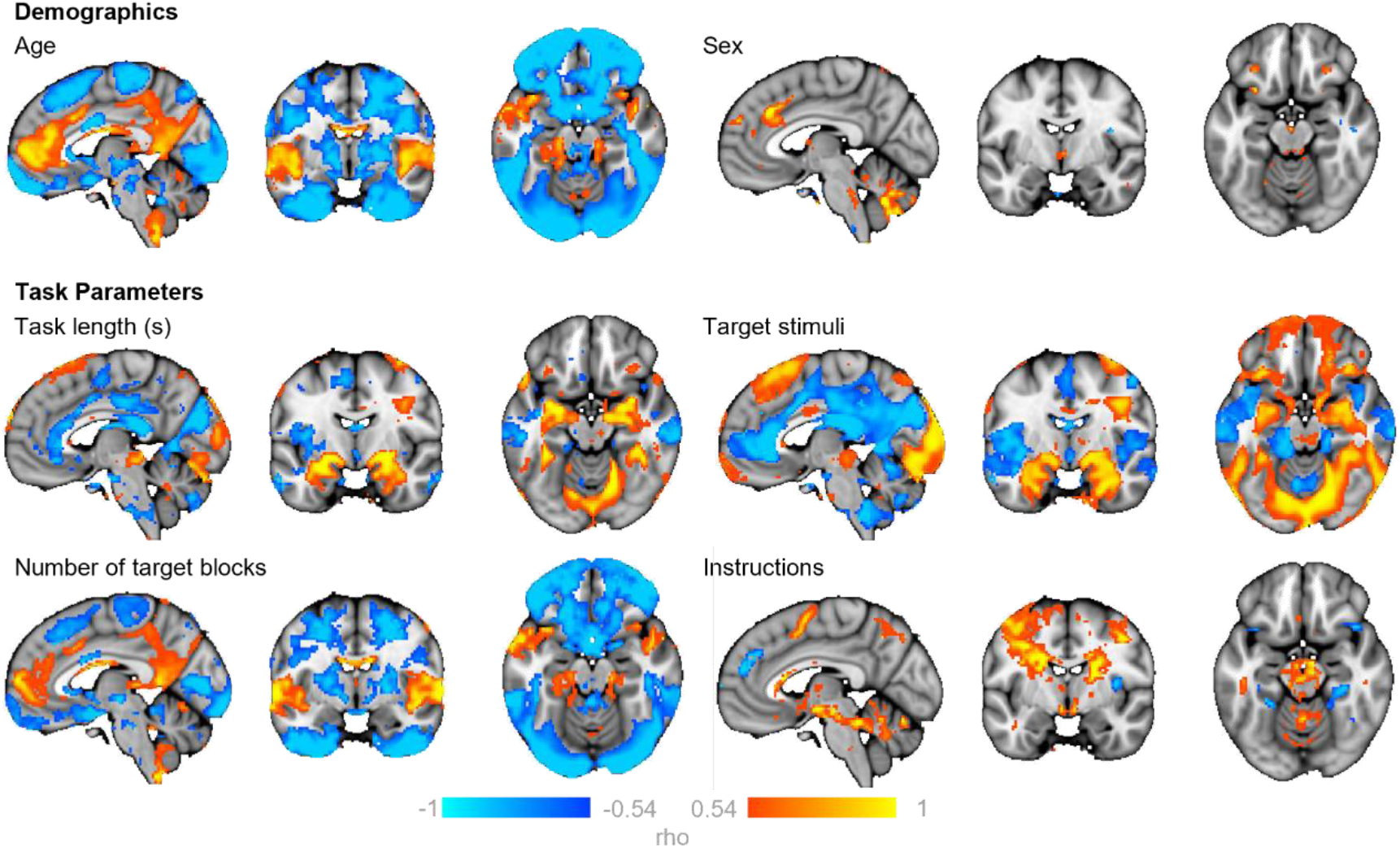
The relationship between input variables and the predicted BOLD activation for faces>shapes. Maps show the correlation coefficients (rho) thresholded by their respective coefficients of determination (rho^2^>0.3) of selected model input variables. This can be interpreted as showing how predicted BOLD activation for the faces>shapes contrast relates to the input variables of the normative models. Positive correlations (warm colours) indicate greater activation for higher values of the input variable and negative correlations (cool colours) greater activation for lower values of the input variable (note that some variables are dummy coded, e.g. target stimuli, instructions). x,y,z = -4,-6,-16.

**Table 1:**
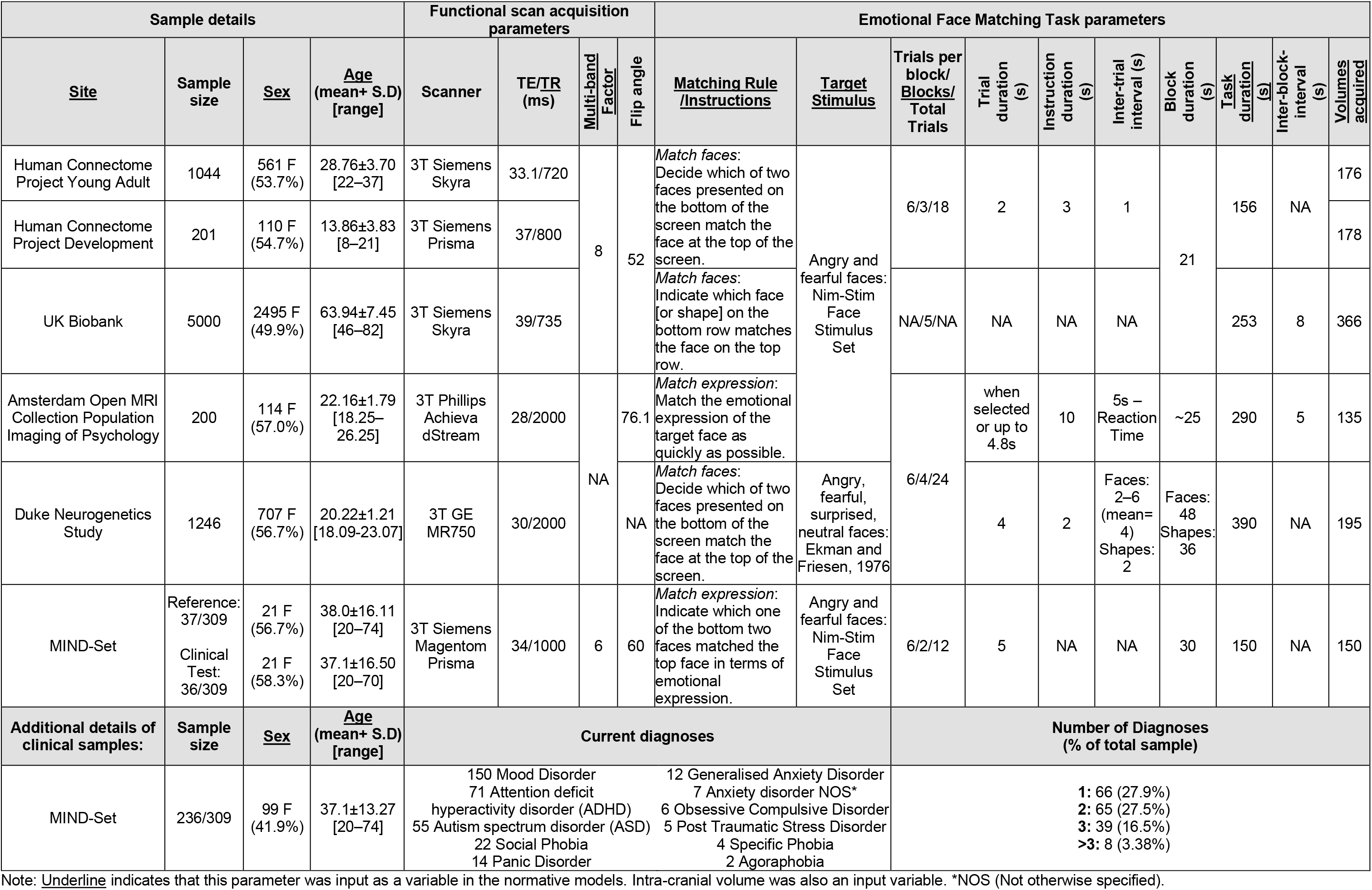
Sample details, functional scan acquisition parameters and Emotional Face Matching Task parameters for data included in the normative models.

### Application of normative model to a naturalistic clinical sample

Next, we aimed to relate the deviations from these normative models to psychopathology. To achieve this, we evaluated the patient cohort with respect to the normative models estimated from the large reference cohort. For the faces>shapes and faces>baseline models, the explained variance of the clinical test data was quite low. This was expected given that this cohort is quite homogenous with respect to the covariates included in the model (i.e., all subjects were scanned on the same scanner, using the same experimental paradigm and had an age range considerably narrower than the reference cohort). This suggests that the variance in BOLD signal was driven more by individual differences, as opposed to the variables included in the model. The skew and kurtosis of the models were centred around zero. See Supplementary Figure 7a-h for histograms of these evaluation metrics, and their respective illustration on the brain.

### Frequency of deviations differentiates patients from reference test cohort

Next, we compared the frequency of extreme deviations (NPMs thesholded at > ±2.6), at the level of each individual, between patients from the MIND-Set cohort and the reference test cohort for each model type (faces>shapes: Fig. 5b,c; faces>baseline: Fig. 5e,f). MIND-Set patients had a greater frequency of deviations relative to the reference test cohort for the faces>shapes contrast (Mann-Whitney U test = 341806.5, *p* = 2.029^-10^; Fig. 5b). These deviations were most frequently identified in the lateral ventral prefrontal cortex, and the bilateral medial and inferior temporal lobe (Fig. 5a). In contrast, for the faces>baseline contrast individuals there was no significant difference in the frequency of deviations between the reference test cohort relative to MIND-Set patients (Mann-Whitney U test = 487921.0, *p* = 0.22; Fig. 5e). These comparisons were repeated excluding the vmPFC region and the main effects changed minimally for both contrasts.

**Figure 5:**
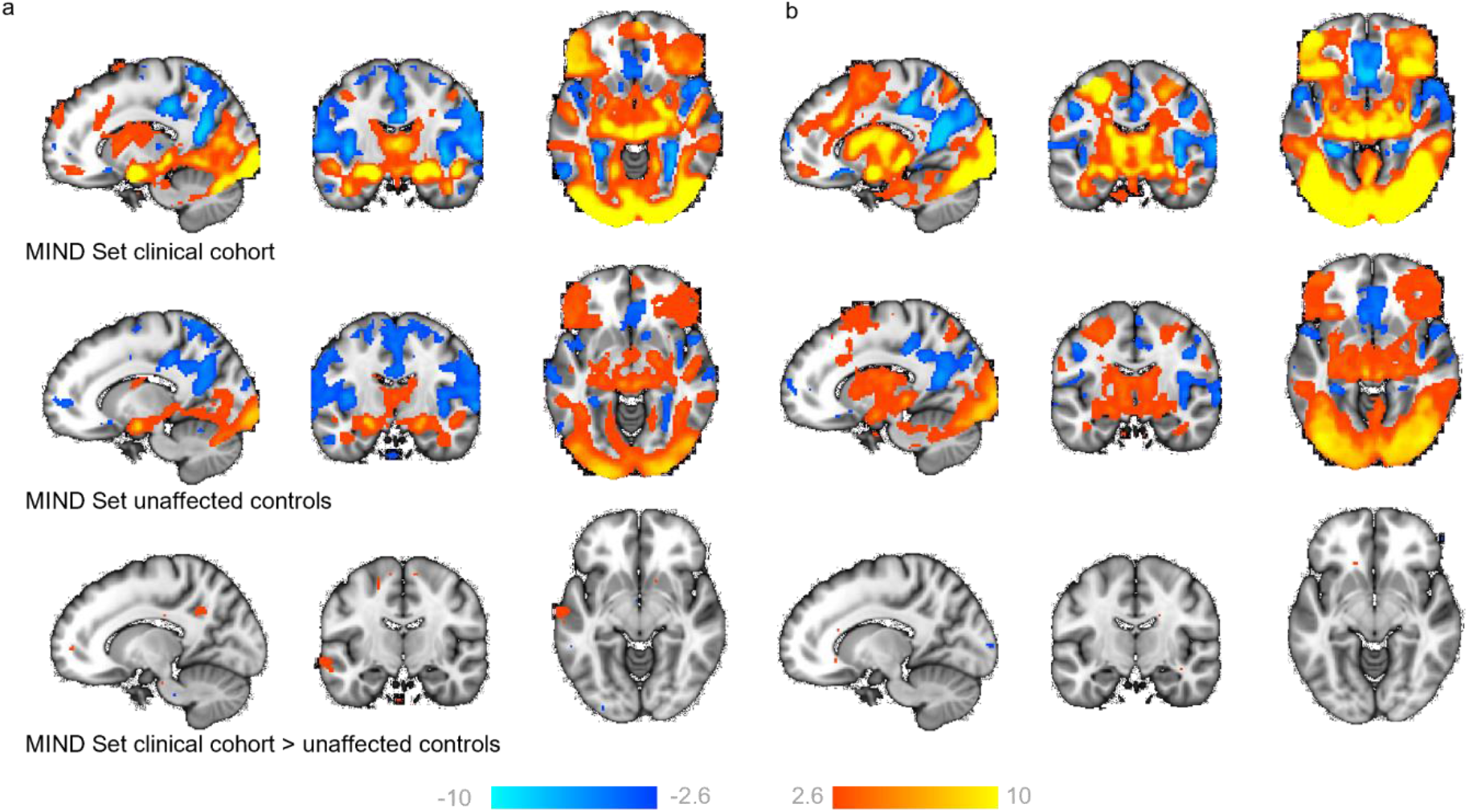
General linear model results comparing patients to controls for the faces>shapes and faces>baseline contrasts. Maps show regions activated (warm colours) and deactivated (cool colours) for faces>shapes (a) and faces>baseline (b), for patients (top row) and unaffected controls (middle row) from the MIND Set cohort. (c) Regions where patients have more activation than controls (bottom row) (z-statistic maps, thresholded at > ±2.6). x,y,z = -14,-13,-9.

### Associations of patterns of deviation with cross-diagnostic symptom domains

We then aimed to determine whether multivariate patterns of deviation from the reference models were associated with cross-diagnostic symptomatology. To achieve this, we input whole-brain deviation maps (unthresholded, such that any deviations, irrespective of their magnitude, could potentially contribute to the observed correlations and removing any risk of bias) and factor loadings for negative valence, cognitive function, social processes and arousal/inhibition domains from prior work^17^ to an established penalised canonical correlation analysis (CCA) framework that enforces sparsity (sparce CCA, SCCA; functional domain loading scores were available for 217 patients)^18,19^. Significant out of sample associations (10 fold 70% - 30% training-test split) were detected both for faces>shapes and faces>baseline contrasts (mean *r* of test splits 0.183 and 0.216 respectively, both *p* < 0.001 by permutation test; Fig. 6b, e) but with distinct patterns of effects both in terms of symptom domains and associated brain regions. More specifically, for the faces>shapes contrast, decreased functioning predominantly in the negative valence and arousal/inhibition domains (Fig. 6a) was associated with a pattern of deviations including the right insula, the bilateral medial prefrontal cortex and pre-and post-central gyri, the bilateral inferior temporal gyrus, lingual gyrus, bilateral hippocampus and the right thalamus, as well as the regions in the medial and left lateral cerebellum (Fig. 6c). By comparison, for the faces>baseline contrast factor loadings for cognitive functioning and arousal/inhibition (Fig. 6d) were most strongly related to a pattern comprising bilateral insula, the anterior-to-medial cingulate cortex extending to the dorsal medial prefrontal cortex, the pre-and post-central gyri, the right middle frontal and bilateral inferior frontal gyrus, and the bilateral hippocampus, caudate, putamen and amygdala, and the medial and left lateral cerebellum (Fig. 6f). The SCCA was repeated to relate participant’s diagnoses with their whole-brain (unthresholded) deviation maps. In contrast to the cross-diagnostic symptom domains, there was no association between diagnostic labels and deviation scores. Mean canonical correlations were small (mean *r* of test splits <0.1 for both faces>shapes and faces>baseline models), and this was not statistically significant as determined by 1000-fold permutation testing. We repeated the SCCA, first using a grey matter mask and then again using a mask of task positive regions, and the results changed minimally (see Supplementary Figure 10).

**Figure 6:**
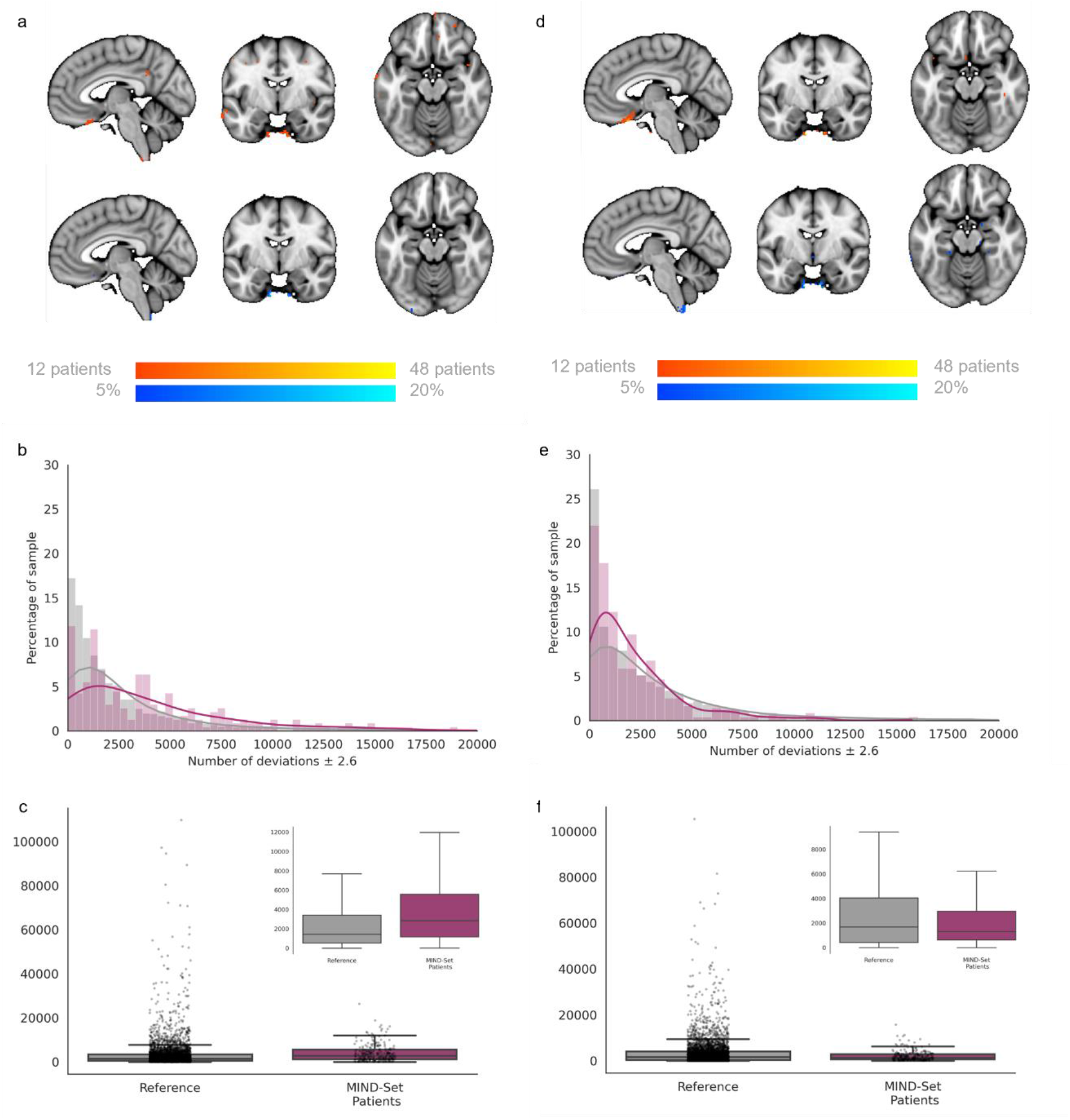
Testing the faces>shapes (left) and faces>baseline normative models with the MIND-Set cohort. Normative Probability Maps illustrate the percentage of participants of the clinical sample who had positive (hot colours) or negative deviations (cool colours) > ±2.6 within each voxel, for the faces>shapes (a) and faces>baseline (d) models. Histograms and box plots show the relative frequency and mean number of the total deviations that a participant has for faces>shapes (b,c) and faces>baseline (e,f) models. x,y,z = -4,-6,-16.

**Figure 7:**
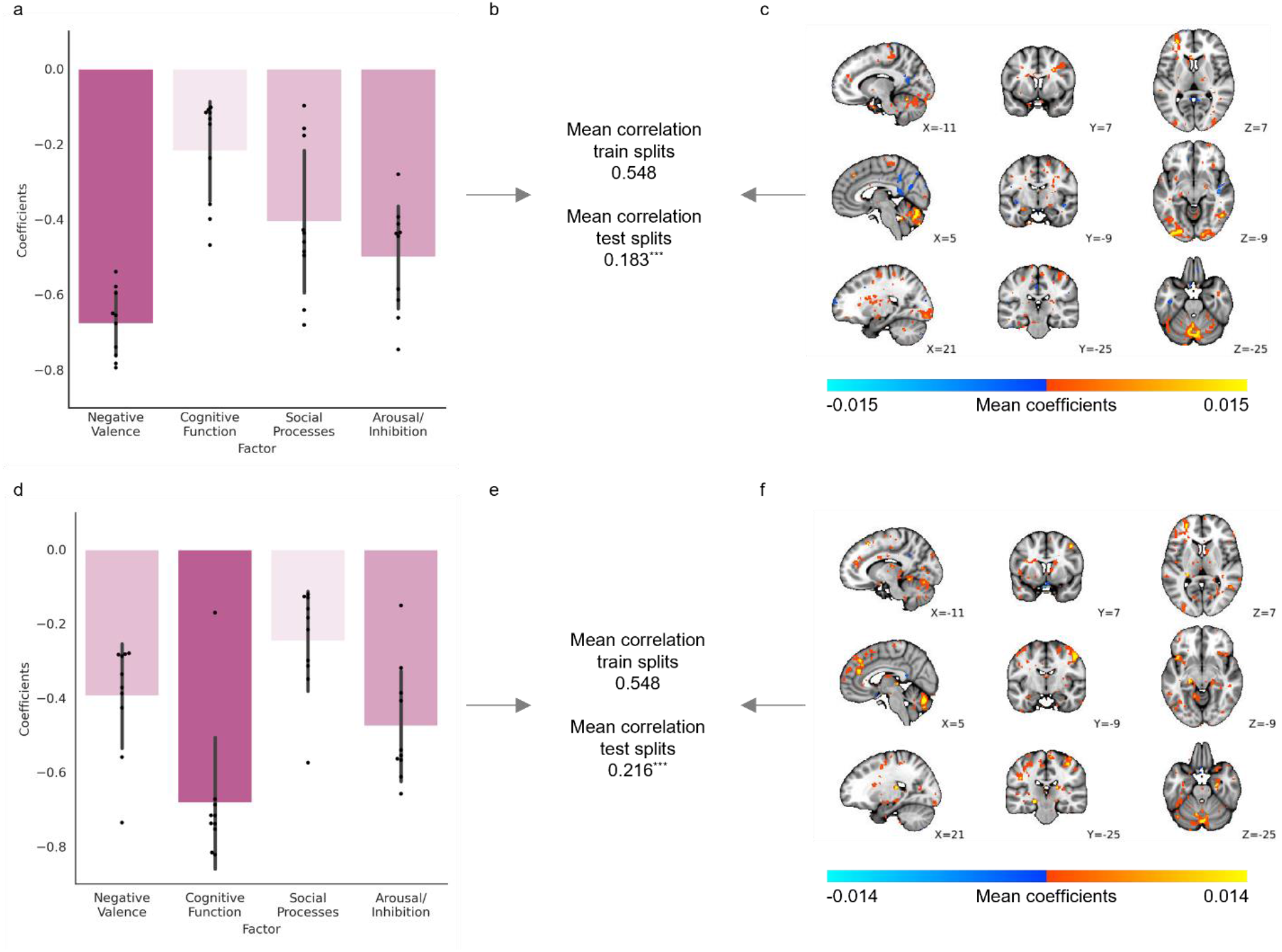
Sparse canonical correlation analyses (SCCA) between functional domains, and deviation scores from faces>shapes or faces>baseline normative models. Weights per factor to latent variable of psycho-social functioning (a,d). Canonical correlation between 4 functional domains and whole-brain deviation scores from (b) faces>shapes and (e) faces>baseline normative models (regularisation 10%). Mean voxel-wise weights to latent variable of deviation scores from (c) faces>shapes normative models and from (f) faces>baseline. All results are statistically significant with 1000-fold permutation tests (*** = *p*<0.001).

### Spatial extent of deviations highlights similarities across, and differences between diagnoses

Finally, we were interested in mapping the spatial distribution of the deviations within the clinical sample, and whether this varied according to the participant’s mental health diagnosis or neurodivergence (note that subjects can be in multiple categories; Sup Fig. 5). For each diagnosis, the pattern of deviations was highly heterogeneous, providing further support for high degree of inter-individual heterogeneity we have reported previously for mental disorders^4,5,7^, and underlining the need to move beyond case-control comparisons at the level of diagnostic groups.

## Discussion

In this study we made use of six large publicly available datasets of participants completing the fMRI EFMT to build a reference normative model of functional activation underlying emotional face processing. We collated data from over 7500 participants and show that our voxel-wise models can explain up to 50% of variance in observed BOLD signal, with the remaining unexplained variance representative of individual differences in functional activation (deviation scores). We unpacked the variance explained by the models, to show how the predicted activation related to the models’ input variables, namely demographics, variations in task design, and acquisition parameters. Lastly, we tested our reference model with data from a sample of patients with heterogenous and frequently co-occurring psychiatric conditions (mood and anxiety disorders, and neurodevelopmental conditions). Our analyses show that: (i) there is considerable inter-individual variation superimposed on the group effects customarily reported in fMRI studies, (ii) that such variation is predictive of psychiatric symptom domains in a cross-diagnostic fashion and (iii) while an overall effect of diagnosis was evident, this was highly individualised in that the overlap of deviations amongst individuals with the same diagnosis was low. This implies that there are brain regions wherein patients more frequently have deviations irrespective of the type of diagnoses, and other regions wherein the frequency of deviations appears specific to the mental-health condition or neurodivergent diagnosis.

A key feature of the normative modelling framework in the context of multi-site fMRI data is that it allows us to aggregate data across multiple samples by binding them to a common reference model. This provides multiple benefits: it removes site effects from the data without requiring the data to be harmonized^20^, which avoids the introduction of certain biases due to harmonisation^21^ and allows meaningful comparisons to be drawn across studies. For example, this allows aggregation of different studies to better understand variation across cohorts or across the lifespan and to understand the effect of different task parameters on functional activity across cohorts. Moreover, by placing each individual within the same reference model this provides the ability to quantify, compare and ultimately parse heterogeneity across studies.

Traditional group-level task contrasts, as shown in Fig. 1, inform us of the region’s most consistently activated across participants/groups during task conditions. Their interpretation has relied heavily on the assumption of spatial homogeneity of activation between subjects; an assumption that the deviation scores from our reference model show to be largely untrue (Fig. 2). We show that such group effects reflect a small proportion of the variation amongst individuals and using the normative modelling framework we map the underlying heterogeneity, separating variation in the intensity and spatial extent of task-evoked functional activation between-subjects attributable to known factors such as site effects, demographics, acquisition parameters, and differences in EFMT paradigm design. More importantly, we show that residual differences in the neuronal effects elicited by the task are highly meaningful in that they were predictive of psychiatric symptomatology and can be used to understand inter-individual differences in functional anatomy and its relation to clinical variables. In our test reference population, while every voxel of the brain had at least one participant with a large deviation, some regions considered active during the faces condition (as compared to shapes), such as the medial occipital lobe, fusiform gyrus and inferior temporal lobe, were also regions in which positive deviations were frequently observed.

When building our reference models, we chose to include and control for multiple variables that we reasoned may influence the BOLD signal observed. These included demographic factors such as age and sex, and task design choices that could influence the BOLD signal generated, as well as acquisition parameters that could influence the BOLD signal recorded. Some effects, such as that of age and task instructions were relatively strong and interpretable, for example, increased age predicted decreased activity in surface areas of the brain and regions surrounding the ventricles likely reflecting decreased signal due to age-related atrophy, and instructing participants to match emotional expressions, as opposed to matching faces, increased the predicted activity in the thalamus which may reflect increased engagement of regions associated with affective processing. On the other hand, other variables explained relatively little variance in the predictions (e.g. sex). In our sample many predictor variables were collinear across sites which limited our ability to detect systematic differences resulting, for example, from differences in the task paradigm or age. In this work we decided to keep all variables in the model and used structure coefficients to identify the importance of different variables, which are relatively insensitive to collinearity. This follows prior work to identify specific effects of input variables on model predictions, for example the influence of specific adversity types on predicted morphometric changes (Holz et al., *in prep*). However future researchers may consider reducing the dimensionality of their inputs prior to model construction. Future studies with larger numbers of more diverse samples (e.g. more variations on the basic task design) that include participants across the entire lifespan, as is possible in consortium such as ENIGMA, will allow for more fine grained analyses of the effect of task parameters on inter-individual variation within the population. Despite our efforts to collect a large representative sample of participants completing an EFMT, participants in mid-adulthood (specifically aged between 37 and 46) were under-represented in our sample. While we do not expect our results to change dramatically with the inclusion of additional participants, as previous studies suggest any age-related changes are best captured by gradual linear or second order trends, future studies should aim to ensure a continuous and overlapping age distribution from samples to minimise the conflation of predictor variables.

We demonstrated that distinct patterns of deviations, derived from each model type (faces>shapes or faces>baseline), were associated with unique profiles of functioning across four transdiagnostic domains. The distinct patterns of effects, in terms of the implicated symptom domains and associated brain regions, make sense in the context of relevant existing literature. For example, negative affect, impulsivity and emotional liability have previously been related to functional activity within the bilateral insula, motor cortex and hippocampus^22^, and cognitive functioning has been linked to activity within the medial prefrontal cortex, anterior-to-medial cingulate cortex, superior frontal gyrus. This not only validates the interpretability of findings from these normative modelling analyse, but also illustrates the potential for future researchers to use individualised deviation maps to better understand the neural processes that underly cognitive and affective functioning, within and across diagnostic boundaries. Furthermore, approaching dysfunction through the normative modelling framework and transdiagnostic functional domains appears to more closely relate to underlying biology. This reflects practitioners implementation of clinical care and the use of overlapping treatments for differing disorders, which often does not fit a binary classification paradigm. Using this modelling approach may also better allow for the quantification of neurodivergence, not as being ‘disordered’ but rather as varying phenotypic expressions along a characterised spectrum.

It should be noted, however, that within any one voxel of the brain, only ∼20% of the clinical sample (be that in the total sample, or within disorders) had large deviations. This suggests that the exact location of deviations is very variable between individuals, and could explain why many prior studies have not found significant differences when performing traditional case-control analyses. In this study, we aimed to estimate the degree to which the deviations from the normative models were associated with cross-diagnostic symptomatology, but other approaches may also be useful, as outlined in Rutherford, et al. ^23^. For example, clustering algorithms could be applied to derive a stratification of individuals^4,5^ or to identify heterogenous yet convergent functional processes (many-to-one functional mappings)^24^, and supervised learning methods may be useful to assess the degree to which specific clinical variables can be predicted from the patterns of deviation we report.

Interestingly, the normative models of functional activation built using the faces>shapes have a different pattern of association with symptomatology relative to the faces>baseline contrast. This suggests that the two contrasts carry complementary information about psychopathology. The frequency of deviation scores was significantly greater in the clinical cohort, compared to the reference cohort, when using the faces>shapes contrast, and the weights attributed to each of these deviations (at a voxel-level) in the SCCA were associated with different symptom domains. Neither contrast was significantly predictive of diagnosis. By comparison, the relationship between the frequency of deviation scores and domains of function was stronger when using models built using the faces>baseline contrast, which was further supported by the stronger canonical correlation between factor loadings for functional domains and deviations from the faces>baseline models. Taken together, this could be interpreted to suggest that widespread deviations, best captured by the faces>shapes contrast, are indicative of global alterations in functioning which are broadly linked to different clinical diagnoses. By comparison, fewer but more focal deviations and/or the ability to detect abnormal baselines of activation^25,26^, best revealed using the faces>baseline, have greater relation to specific functional domains. Future researchers should carefully consider the task contrast used to construct their normative models.

Concerns for the within-subject reliability of task-based fMRI data^27^ are not to be dismissed in the context of our models which are currently built on cross-sectional data. While we acknowledge the limitations imposed due to the general limits of test-retest reliability of task fMRI, our results encouragingly show that the deviation scores from a normative model appear replicable over Test-Retest scans. Comparisons between individuals’ voxel-wise deviations suggested that the magnitude of scores remained stable across ∼95% of the brain, and further identified regions where deviation scores were most influenced by session (i.e. vmPFC regions likely due to session specific signal drop out). The normative modelling method further provides encouraging evidence for the use of task fMRI readouts as individualised biomarkers as we show by their ability to predict clinical variables in the context of SCCA. The normative modelling framework is also ideally positioned to directly test the reproducibility of fMRI within subjects. In follow-on work to the present manuscript, we are currently developing an extension to explicitly include test-retest variability in the model by testing reference models with repeat scans from participants, and compare individuals’ deviation scores between the two tests, whilst explicitly quantifying within subject variance, such that it provides a lower bound on the size of deviation that can be considered meaningful (Bučková et. al., *in prep*). Alternatively, where multiple repeats are available, hierarchical models can be used to accommodate dependencies between subjects^20^ which would provide more precise estimates of individual deviations. The application of the normative modelling method to fMRI can easily be generalised to other tasks (e.g. the monetary incentive delay incentive processing task or n-back work memory task) and need not stop at predicting functional activation. With the right data sets, this method could use fMRI data to predict many other variables including psychophysiological responses or subjective ratings of affect.

With this work, we show the potential for the normative modelling framework to be applied to large task-based fMRI data sets to bind heterogeneous datasets to a common reference model and enable meaningful comparisons between them. Using this approach, we illustrate the heterogeneous intensity and spatial location, the reliability of task-evoked activation within the general population^2^ using the EFMT in a sample of over 7500 participants. Further, we applied this model to patients with a current diagnosis (mood and anxiety disorders, ASD and/or ADHD) and demonstrate the transdiagnostic clinical relevance and further potential for deviation scores derived from this method. The potential of this method is clear; normative modelling of task-based functional activation can facilitate a better understanding of individual differences in complex brain-behaviour relationships, and further our understanding of how these differences relate to mental health and neurodivergence.

## Methods

### Data sets

We collated a large reference sample from 6 independent sites for whom high quality fMRI data for the EFMT are available: AOMIC PIOP2, Duke Neurogenetics Study, HCP Development, HCP Young Adult (1200 release), UK Biobank, and the MIND-Set cohort which also includes a clinical population. For sample details per site see Table 1. Informed consent was obtained from all participants, and, for publicly available datasets ethical approval was provided by the relevant local research authorities for the studies contributing data. The MIND-Set study was approved by the Commissie Mensgebonden Onderzoek Arnhem-Nijmegen.

### fMRI task paradigms

All sites collected a variant of the EFMT^9^. Although specific parameters varied, the overall design was consistent: in each face trial participants were presented with three images of human faces in a triangular formation. Participants were instructed to identify which of two faces/expressions presented at the bottom of the screen matched the one presented at the top of the screen by pushing a button with the index finger of their left or right hand. Multiple face trials were presented in one face block, and the task included multiple face blocks (see Table 1 for the number of trials per block, and number of blocks per site). As a somatomotor control, participants also completed shape trials, wherein they were presented with three geometric shapes (circles and ovals) and asked to indicate which of the two shapes presented at the bottom of the screen matched the one at the top. Multiple shape trials were concatenated to form one shape block, which were interspersed between face blocks. For further information about the control stimuli, see Supplementary Table 1).

Two paradigms (HCP Young Adult and HCP Development) included an inter-trial interval (white fixation cross on black screen), and three sites (HCP Young Adult, HCP Development and AOMIC PIOP2) had an instruction trial that preceded the start of each block. Tasks varied in their duration from 150 to 290 seconds, which indirectly corresponded to the acquisition of between 135 and 336 functional volumes.

#### fMRI data acquisition

Site specific acquisition parameters per site are detailed in Table 1, and in the following site specific protocols: AOMIC PIOP2^15^, HCP Young Adult^13^, HCP Development^28^, UKBiobank^29^, Duke Neurogenetics Study (https://www.haririlab.com/methods/amygdala.html) and MIND-Set^30^.

#### fMRI pre-processing

Data pre-processing was harmonised across all sites; a FSL-based pipeline^31^ was consistently applied to decrease the likelihood of introducing variance due to pre-processing differences. Since the HCP young adult, HCP development and UKB Biobank data were already processed relatively consistently, we reused the processing pipelines provided by the respective consortia (for HCP sites we used the minimal processing pipeline)^29,32^, with additional steps taken as necessary (e.g. matching smoothing kernels across studies). At a within-subject level, all functional data underwent gradient unwarping, motion correction, fieldmap-based EPI distortion correction (where fieldmaps were available), boundary-based registration of EPI to structural T1-weighted scan, denoising for secondary head motion-related artifacts using automatic noise selection, as implemented in ICA-AROMA^33^, non-linear registration into MNI152 space, and grand-mean intensity normalization. Where applicable, datasets were resampled to 2mm^3^ resolution, and all data were spatially smoothed using a 5 mm FWHM Gaussian kernel.

#### Quality control

Participants were excluded if their mean relative RMS was greater than 0.5mm. Additional quality control was performed for signal coverage in the prefrontal cortex for the UK Biobank sample (see supplementary methods).

#### fMRI general linear modelling (GLM) – single subject

We matched the methodological approach used to estimate the parameters within a GLM-based analysis, given evidence to suggest this analytic step can significantly contribute to the variability of reported results between sites^34^. Therefore, for each site, the linear model parameter were estimated using the FSL software package version 6.03 (HCP Young Adult, HCP Development, MIND-Set, Duke Neurogenetics Study; http://fsl.fmrib.ox.ac.uk/) or as downloadable form UK Biobank^29^. Two regressors were constructed from the faces and shapes blocks which were then convolved with a canonical double-gamma haemodynamic response function and combined with the temporal derivatives of each main regressor. These were treated as nuisance regressors and served to accommodate slight variations in slice timing or in the haemodynamic response. Data were pre-whitened using a version of FSL-FILM customized to accommodate surface data, the model and data were high-pass filtered (200s cut-off). Fixed-effects GLMs were estimated using FSL-FLAME 1: first for independent runs, then when necessary combining two runs into a single model for each participant (HCP Young Adult). and the AOMIC, DNS and MIND-Set maps were transformed into standard space using FNIRT^35^. We created summary group level maps per site (for a random sample of 100 participants; see Figure 1 and Supplementary Figure 3), as a sanity check to ensure the data was otherwise comparable to past literature and performed a case-control comparison between patients with a current diagnosis (mood and anxiety disorders, ASD and/or ADHD) and unaffected controls in the MIND-Set cohort (see Figure 5).

#### Normative models

The z-statistic maps from the contrast face>shapes (5mm smoothed in standard space), for each subject, were used as response variables for the normative models. That is, we specified a functional relationship between a vector of covariates and responses. We created normative models of EFMT-related BOLD activation maps, as a function of site, age, sex, intra-cranial volume, and acquisition [TR, multiband sequence, number of volumes collected (per run where applicable)] and task parameters [task length (in seconds, and per run where applicable), number of target blocks, instructions for faces condition, and the target stimuli (i.e. which stimulus set they came from)], by training a Bayesian Linear Regression (BLR) model to predict BOLD signal for the faces>shapes contrast. Generalisability was assessed by using a half-split train-test sample (train: 3885, test: 3843). In preliminary analyses, we compared a warped model which can model non-Gaussianity with a vanilla Gaussian BLR model. Since the fit was comparable across most metrics and regions, we focus on the simpler Gaussian model below. We included dummy coded site-related variables as additional covariates of no-interest. We also created models to predict BOLD signal for the faces condition alone (i.e. face>baseline contrast; train: 3778, test: 3950 split). This was performed in the Predictive Clinical Neuroscience toolkit (PCNtoolkit) software v0.26 (https://pcntoolkit.readthedocs.io/en/latest) implemented in python 3.8.

#### Quantifying voxel-wise deviations from the reference normative model

To estimate a pattern of regional deviations from typical brain function for each participant, we derived a normative probability map (NPM) that quantifies the voxel-wise deviation from the normative model. The subject-specific *Z*-score indicates the difference between the predicted activation and true activation scaled by the prediction variance. We thresholded participant’s NPM at Z = ±2.6 (i.e. *p* < .005)^7^ using fslmaths and summed the number of significantly deviating voxels for each participant, and then across the total sample.

#### Test-Retest reliability of voxel-wise deviation scores

To determine the Test-Retest reliability of the voxel-wise deviations, we utilised the HCP Retest data (n = 42, mean age = 30.21±3.43 years, 14F). HCP Retest data was pre-processed as detailed above for the HCP Young Adult site. We re-generated the aforementioned normative models removing all participants from the original HCP Young Adult sample for whom Retest data was available. We then tested the new normative models with the original HCP Young Adult (Test) data and the Retest data; i.e. participant 1-n Test scans, and Participant 1-n Retest scans were independently tested against the new normative models. We determined the change in deviation score per voxel, both within-and across-subjects (mean difference) using paired t-test (scipy.stats). We determined the Pearson correlation coefficient between voxel-wise deviations from the Test vs Retest data.

#### Supplementary out of sample test of reference normative models

We collated a new sample of 5000 participants from UK Biobank to test against the reference models. These were the next 5000 participants from the UK Biobank population (2325 F; mean age 63.42±7.54 years), as ranked by vmPFC coverage (i.e. decreasing data quality in this region). Results of this analysis replicated the main findings and can be found in the Supplementary materials.

#### Effects of input variables on model predictions

In order to probe the magnitude of the influence of task design parameters on the predicted BOLD signal, we examined the structure coefficients (correlation coefficients) of each input variable. This approach is preferable to regression coefficients when variables are collinear^36^. Selected structure coefficient maps are displayed.

#### Clinical application

We tested the normative models we created using the reference data, with a heterogeneous patient sample from the MIND-Set cohort (n = 236, mean age = 37. 1±13.27; 41.94% female). This is a naturalistic and highly co-morbid sample derived from out-patients of the psychiatry department of Radboud University Medical Centre. This included 150 patients diagnosed with a current mood disorder (unipolar or bipolar depressive disorder), 12 with generalised anxiety disorder, 22 with social phobia, 14 with panic disorder, 71 with attention-deficit-hyperactive-disorder, and 55 autistic individuals (see Table 1 for full details and note that subjects can be in multiple diagnostic categories). The clinical relevance of our models can also be tested in the context of transdiagnostic symptom domains; a conceptualisation of mental functioning that transcends diagnostic boundaries and allows for nuanced brain-behaviour interpretations. As such, for 217 (of our 236) patients for whom all required data was available, we repeated a previously validated factor analysis method (performed in SPSS v24.0, oblique rotation)^17^ to obtain individual factor loadings on 4 functional domains: (1) negative valence, (2) cognitive function, (3) social processes and (4) arousal/inhibition.

#### Quantifying patients’ voxel-wise deviations from the reference normative model

As for the reference cohort, we generated NPMs to estimate the pattern of regional deviations from typical brain function for each participant, and summed across the sample. We then used a Mann-Whitney U test to compare the frequency of deviations (>±2.6) between the reference controls and patients from the MIND-Set cohort.

#### Relating deviations to transdiagnostic functional domains

In order to map the association of the deviation scores with cross-diagnostic symptomatology, we performed sparse canonical correlation analyses (SCCA) to relate participant’s scores in the four aforementioned functional domains or their diagnoses, to their whole-brain (unthresholded) deviation maps using an established penalised CCA framework that enforces sparsity^18,19^. Specifically, we applied variable shrinkage by adding an l1-norm penalty term to stabilise the CCA estimation and ensure the weights for the deviation scores were more interpretable. We follow the formulation outlined in Witten, et al. ^18^, where we refer to for details. In brief, given two data matrices ***X*** and ***Y*** with dimensions *n* × *p* and *n* × *q* respectively (here, these are the cross-diagnostic factor loadings and whole-brain deviations), and two weight vectors ***u*** and ***ν*** this involves maximising the quantity *ρ* = ***u***^***T***^***X***^***T***^***Yν*** subject to the constraints 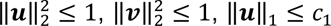 and ‖*ν*‖_1_ ≤ *c*_2_, where the penalties *p*(***u***) and *p*(***ν***) are the standard L1-norm. We set the regularisation parameters for each view heuristically (*c*_1_ = 0.9*p* corresponding to light regularisation for the factor scores, *c*_1_ = 0.1*q*, corresponding to heavy regularisation for the deviation maps such that no more than 10% of voxels were selected). While it is possible that better performance would be obtained by optimising the regularisation parameters across a grid, we did not pursue that here due to the moderate sample size for the clinical dataset. We assessed generalisability of SCCA by splitting the data in to 70% training data and 30% test 10 times. Finally, we wrapped the entire procedure in a permutation test where we randomly permuted the rows of the behavioural matrices 1000 times to compute an empirical null distribution for significance testing. We performed an additional set of sensitivity analyses to more carefully evaluate the sparse canonical correlation analyses (SCCA). Specifically, we repeated the SCCA, first using a grey matter mask (Harvard Oxford probability thresholded at 30% probability), and then again using a mask of task positive regions (HCP Young Adult group level T-statistic map thresholded at t>3.6).

#### Spatial patterns of deviations by primary and co-occurring diagnoses

We illustrated the spatial heterogeneity in deviations between different diagnoses (note that subjects can be in multiple categories), and further, compared patients with a single diagnoses to those with two, three, or more than three diagnoses, to determine whether and if so, how the location of deviations related to the number of co-occurring diagnoses a patient has.

## Data availability

Scripts for running the analysis and visualizations are available on GitHub (https://github.com/predictive-clinical-neuroscience/EFMT_Norm_Models). Group level quality metrics (explained variance, skew, kurtosis, and SMSE) and group level results (frequency of deviations) are also available as nifti files.

## Acknowledgements

The Duke Neurogenetics Study was supported by Duke University and National Institutes of Health (NIH) grants R01DA031579 and R01DA033369. Ahmad R. Hariri (Duke University) is further supported by NIH grant R01AG049789. The Duke Brain Imaging and Analysis Center’s computing cluster, upon which all DNS analyses heavily rely, was supported by the Office of the Director, NIH, under Award Number S10OD021480. CFB gratefully acknowledges funding from the Wellcome Trust Collaborative Award in Science 215573/Z/19/Z and the Netherlands Organization for Scientific Research Vici Grant No. 17854 and NWO-CAS Grant No. 012-200-013.

## SUPPLEMENTARY MATERIALS

### SUPPLEMENTARY METHODS – *Sample Details*

**Supplementary Figure 1:**
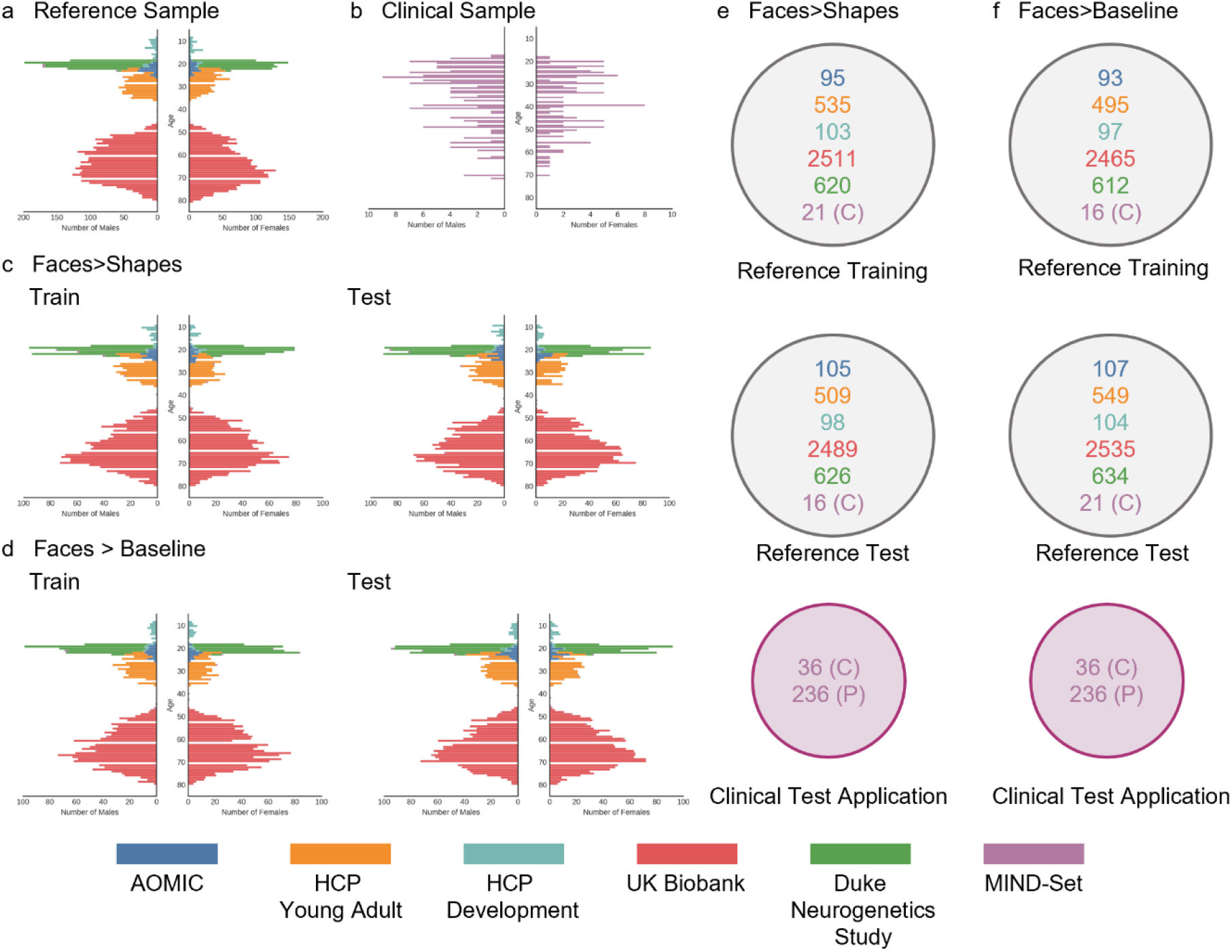
Age and sex distributions and sample splits. Age and sex distributions of (a) the total reference sample, (b) the total clinical test sample (MIND-Set) (c) the faces>shapes train (left) and test (right) split, and (d) the faces>baseline train (left) and test (right) split. Details of the number of participants into the Reference model training and test split, and the clinical test application for (c) faces>shapes (e) and faces>baseline (f). C = unaffected MIND-Set control. P = MIND-Set patient.

### SUPPLEMENTARY METHODS – *fMRI task paradigms*

**Supplementary Table 1:** Additional details of control condition.

| Site | Control Stimulus | Trials per block/<br>Blocks/<br>Total Trials | Trial duration<br>(s) |
| --- | --- | --- | --- |
| Human Connectome Project Young Adult | White outline of circles and ovals presented on black background | 6/3/18 | 2 |
| Human Connectome Project Development |  |  |  |
| UK Biobank |  | NA/5/NA | NA |
| Amsterdam Open MRI Collection Population Imaging of Psychology | Oval stimuli created by pixelating the face stimuli presented on a grey background | 6/5/30 | when selected or up to 4.8s |
| Duke Neurogenetics Study | Black filled circles and ovals presented on a white background |  | 4 |
| MIND-Set | Oval stimuli created by pixelating the face stimuli presented on a white background | 6/2/12 | 5 |

### SUPPLEMENTARY METHODS – *Signal coverage of the prefrontal cortex (PFC)*

Due to air-tissue inhomogeneities which can diminish the acquired BOLD signal to such a degree that no activations are visible, a notorious effect within the ventral PFC, we performed targeted quality control for this extended region. Binary ROI masks were created for the dorsal ventro-medial PFC (d-vmPFC), ventral vmPFC (v-vmPFC), lateral vmPFC (l-vmPFC) and the dorso-medial PFC (dmPFC), as defined by the Harvard Oxford Atlas at 25% probability for atlas regions 25, 27, 33 and 1 respectively (see Supplementary Figure 3B). The percentage of voxels with an absolute value greater than 0 for the contrast faces > shapes within each ROI was determined (i.e. where any signal was present regardless of its relative direction; see Supplementary Figure 1A,B). While most sites had good coverage, the coverage within the ventral and lateral vmPFC regions were particularly variable for the UK Biobank data. We therefore performed this step *only on data from the UK Biobank site*; this selectivity was made possible by the large number of participants we had access to, and our need to include but a fraction of the total available sample. We ranked participants in descending order of the percent of their v-vmPFC, l-vmPFC, d-vmPFC, and dmPFC covered, respectively, and selected the first 5000 participants. We also collected the percentage covered value for a bilateral amygdala ROI mask, but made no exclusion/inclusions on this basis as coverage was very high across all participants and all sites.

**Supplementary Table 2:** Motion QC.

| Site | Full sample size | Included | Excluded |
| --- | --- | --- | --- |
| AOMIC | 217 | 217 | 0 |
| DNS | 1263 | 1246 | 17 |
| HCP Young Adult | 1044 | 1044 | 0 |
| HCP Development | 256 | 256 | 0 |
| UK Biobank | 26167 | 5000 | N/A |
| MIND-Set | 393 | 389 | 4 |

**Supplementary Table 3:** vmPFC QC.

| . | Site | Sample size (n) | Mean percentage of ROI covered | Standard deviation |
| --- | --- | --- | --- | --- |
| Bilateral Amygdala | AOMIC | 217 | 100 | 0 |
|  | DNS | 1246 | 100 | 0 |
|  | HCP Development | 256 | 99.94936 | 0.162845 |
|  | HCP Young Adult | 1044 | 99.98188 | 0.098589 |
|  | MIND Set | 389 | 99.99966 | 0.006707 |
|  | UK Biobank | 26120 | 99.47363 | 2.399575 |
| dmPFC | AOMIC | 217 | 99.10345 | 0.830591 |
|  | DNS | 1246 | 99.61285 | 0.411399 |
|  | HCP Development | 256 | 94.6524 | 2.460686 |
|  | HCP Young Adult | 1044 | 96.85686 | 3.318827 |
|  | MIND Set | 389 | 99.069 | 1.785034 |
|  | UK Biobank | 26120 | 99.0493 | 1.405906 |
| Dorsal vmPFC | AOMIC | 217 | 85.08059 | 12.23881 |
|  | DNS | 1246 | 94.68589 | 4.406996 |
|  | HCP Development | 256 | 72.90154 | 16.11571 |
|  | HCP Young Adult | 1044 | 98.58104 | 2.192929 |
|  | MIND Set | 389 | 99.75218 | 0.886996 |
|  | UK Biobank | 26120 | 94.50698 | 8.107753 |
| Lateral vmPFC | AOMIC | 217 | 98.90096 | 1.063159 |
|  | DNS | 1246 | 99.82392 | 0.33135 |
|  | HCP Development | 256 | 99.14376 | 1.003451 |
|  | HCP Young Adult | 1044 | 99.46709 | 0.978404 |
|  | MIND Set | 389 | 98.23297 | 1.745961 |
|  | UK Biobank | 26120 | 89.56122 | 4.919904 |
| Ventral vmPFC | AOMIC | 217 | 96.06479 | 3.960334 |
|  | DNS | 1246 | 99.45099 | 0.964661 |
|  | HCP Development | 256 | 95.999 | 4.42593 |
|  | HCP Young Adult | 1044 | 99.63665 | 0.85699 |
|  | MIND Set | 389 | 98.83508 | 2.868568 |
|  | UK Biobank | 26120 | 61.17452 | 15.9091 |

**Supplementary Figure 2:**
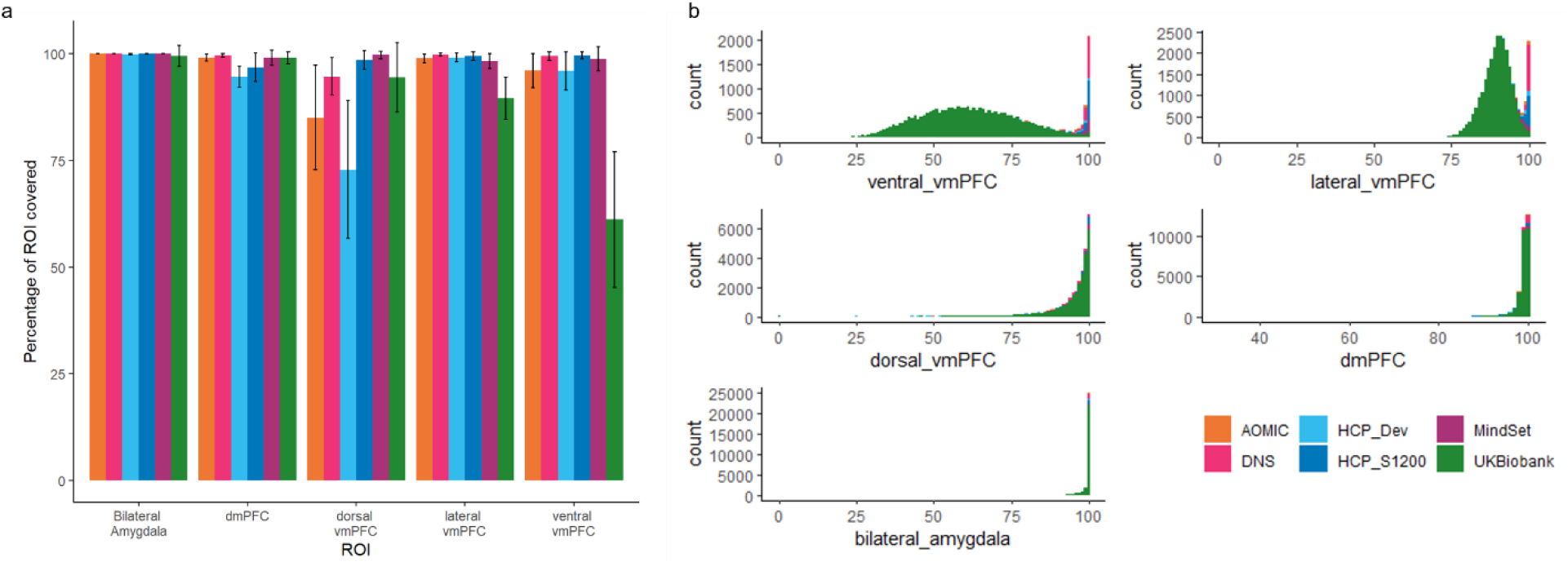
vmPFC QC metrics. (a) Mean percentage of each ROI with signal greater than 0, used for quality control. Error bars show +/- standard deviation (b) Stacked histograms (raw participant count) of the percentage of each ROI covered, coloured by site.

### SUPPLEMENTARY RESULTS – *Overlap of voxels with low tSNR and the prefrontal cortex QC mask*

**Supplementary Figure 3:**
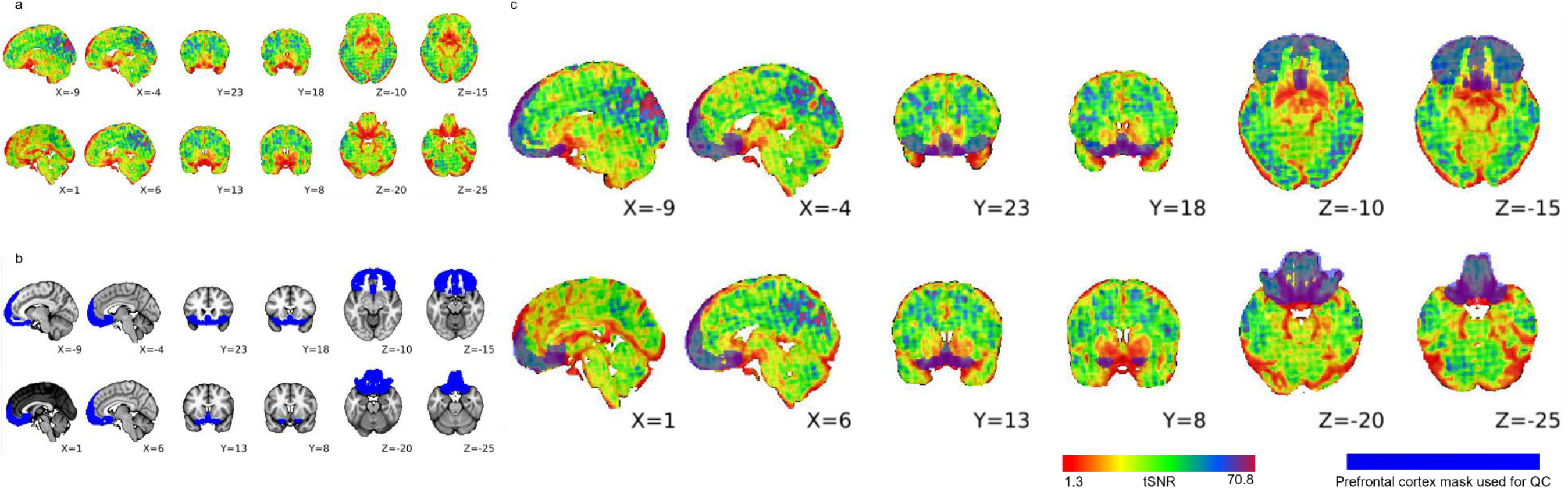
The prefrontal cortex mask used for QC overlaps with regions most affected by low tSNR. A representative example UK Biobank participant’s tSNR map (a) illustrating regions where tSNR is low. (b) The combined prefrontal cortex mask including the dorsal ventro-medial PFC, ventral vmPFC, lateral vmPFC, and the dorso-medial PFC, as defined by the Harvard Oxford Atlas at 25% probability for atlas regions 25, 27, 33. (c) Overlay of mask on tSNR map.

### SUPPLEMENTARY RESULTS – *Task evoked activation*

**Supplementary Figure 3:**
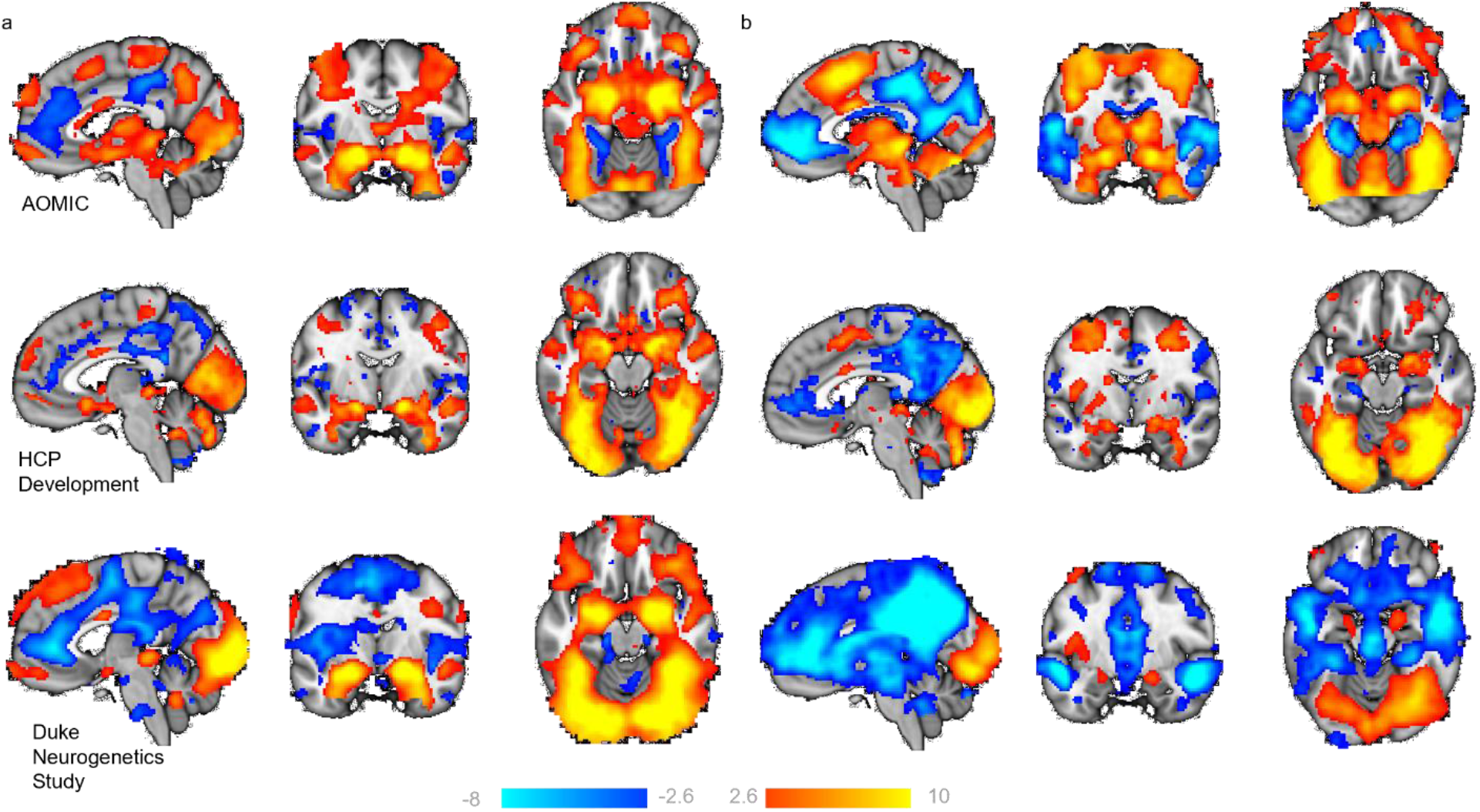
Task evoked activation. *R*epresentative groups maps (from Amsterdam Open MRI Collection Population Imaging of Psychology, HCP Development and Duke Neurogenetics Study), illustrating regions where participants show greater BOLD signal (z-statistic maps, thresholded at>±2.6) to (a) faces, as compared to shapes (faces>shapes), and (b) faces, as compared to baseline (faces>baseline). x,y,z = -4,-6,-16.

### SUPPLEMENTARY RESULTS – *Evaluation of reference normative models*

**Supplementary Figure 4:**
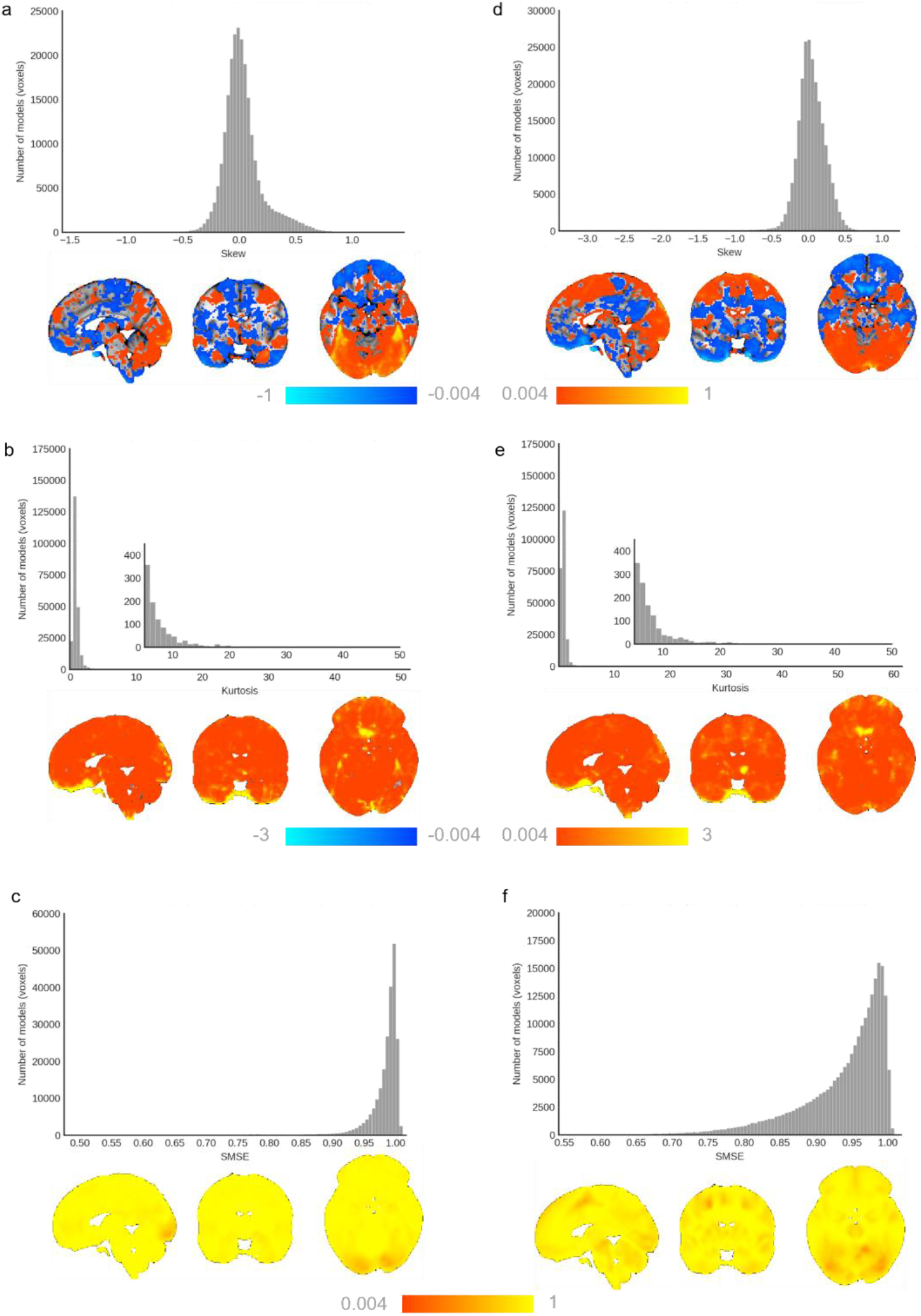
Evaluation of the faces>shapes (left) and faces>baseline (right) reference normative models. Histograms show the skew (a,d), kurtosis (b,e), and SMSE (c,f) of the normative models, and their respective illustration on the brain. x,y,z= 4,-6,-16.

### SUPPLEMENTARY RESULTS – Test-Retest analysis for faces>baseline

**Supplementary Figure 5.**
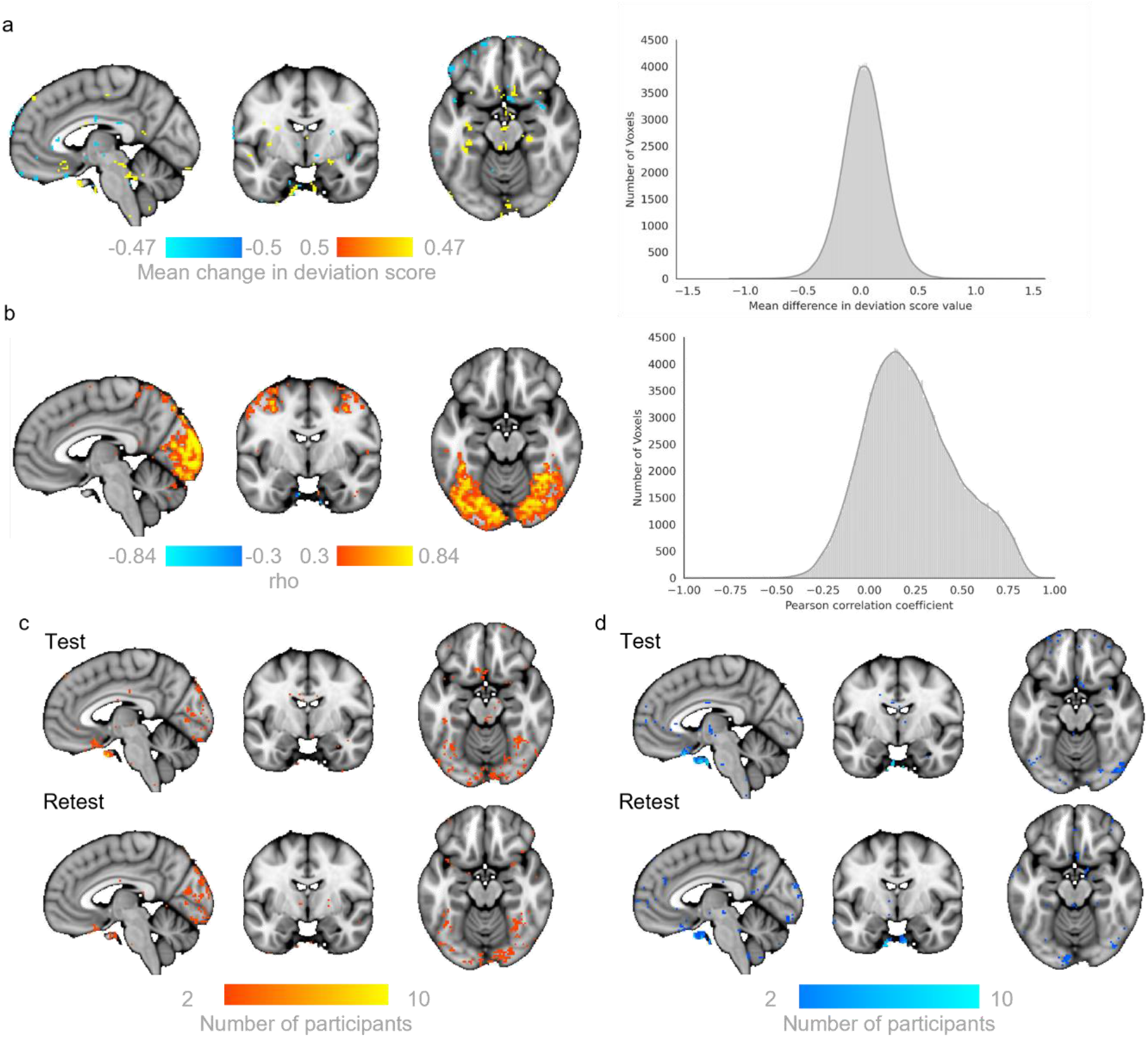
Test – Retest reliability of deviation scores for faces>baseline models. Mean within subject difference per voxel (histogram) illustrated thresholded at >0.5 (i.e. a change greater than half a standard deviation between Test and Retest scans (a). The correlation coefficients (rho) between Test and Retest deviation scores (histogram) illustrated thresholded by the coefficients of determination (rho2>0.3, b). Normative Probability Maps illustrate the voxels wherein 2 or more participant had positive (c, hot colours) or negative deviations (d, cool colours) > ±2.6 for the faces > baseline normative models in the Test (top rows) and Re-Test (bottom rows) samples. x,y,z = -4, -6, -16.

### SUPPLEMENTARY RESULTS – *Additional out of sample test of reference normative models*

We further validated our reference model with an additional out-of-sample/unseen 5000 participants from UK Biobank. These were the next 5000 participants from the UK Biobank population, as ranked by vmPFC coverage (i.e. decreasing data quality in this region). In most brain regions, the results are very comparable to the initial test sample. This includes good explained variance particularly with visual and extrastriate cortex, and subcortical regions including the amygdala; minimal skew and kurtosis; and frequent large positive deviations in visual and extrastriate cortex regions. Relatedly, we note that the results from the models in the vmPFC region show much greater skew and kurtosis than the original sample, and participants more frequently have large negative deviations in this region. Taken together this most likely reflects the worse signal quality in this region and reinforces the relevant influence of data quality on the deviation scores in this area.

**Supplementary Figure 6:**
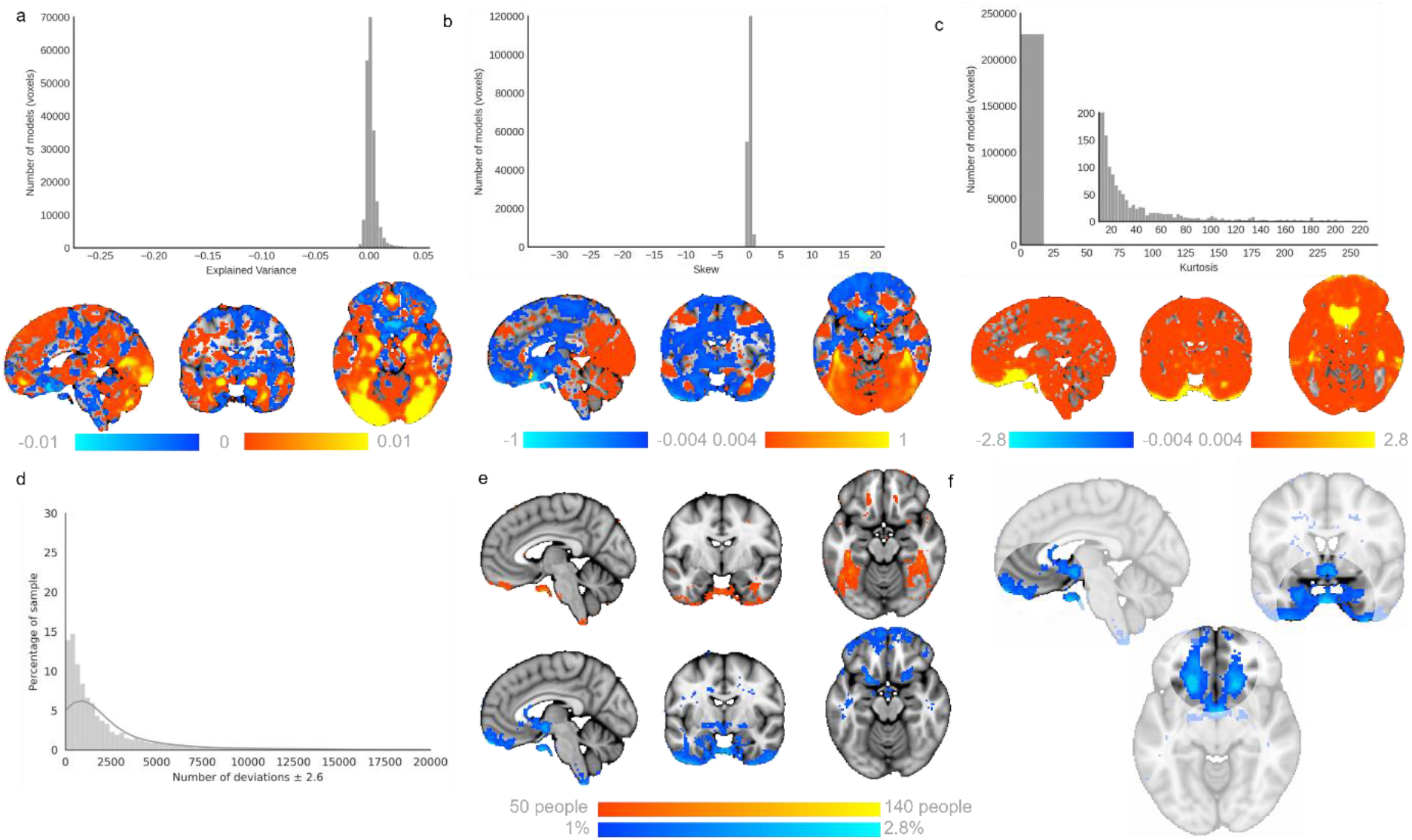
Validation of reference normative models in a new sample of 5000 UK Biobank participants. (a) Explained variance, (b) skew, and (c) kurtosis of results when 5000 new UK Biobank participants were tested on the original reference model. (d) Histogram shows the relative frequency of the total number of deviations that a participant has for each model. (e) Normative Probability Maps illustrate the percentage of participants of the total sample who had positive (hot colours) or negative deviations (cool colours) > ±2.6 within each voxel, for the faces > shapes, where (f) highlights the frequent negative deviations within the vmPFC/OFC region. x,y,z = -4,-6,-16

### SUPPLEMENTARY RESULTS – *Evaluation of normative models when applied to MIND-Set cohort*

**Supplementary Figure 7:**
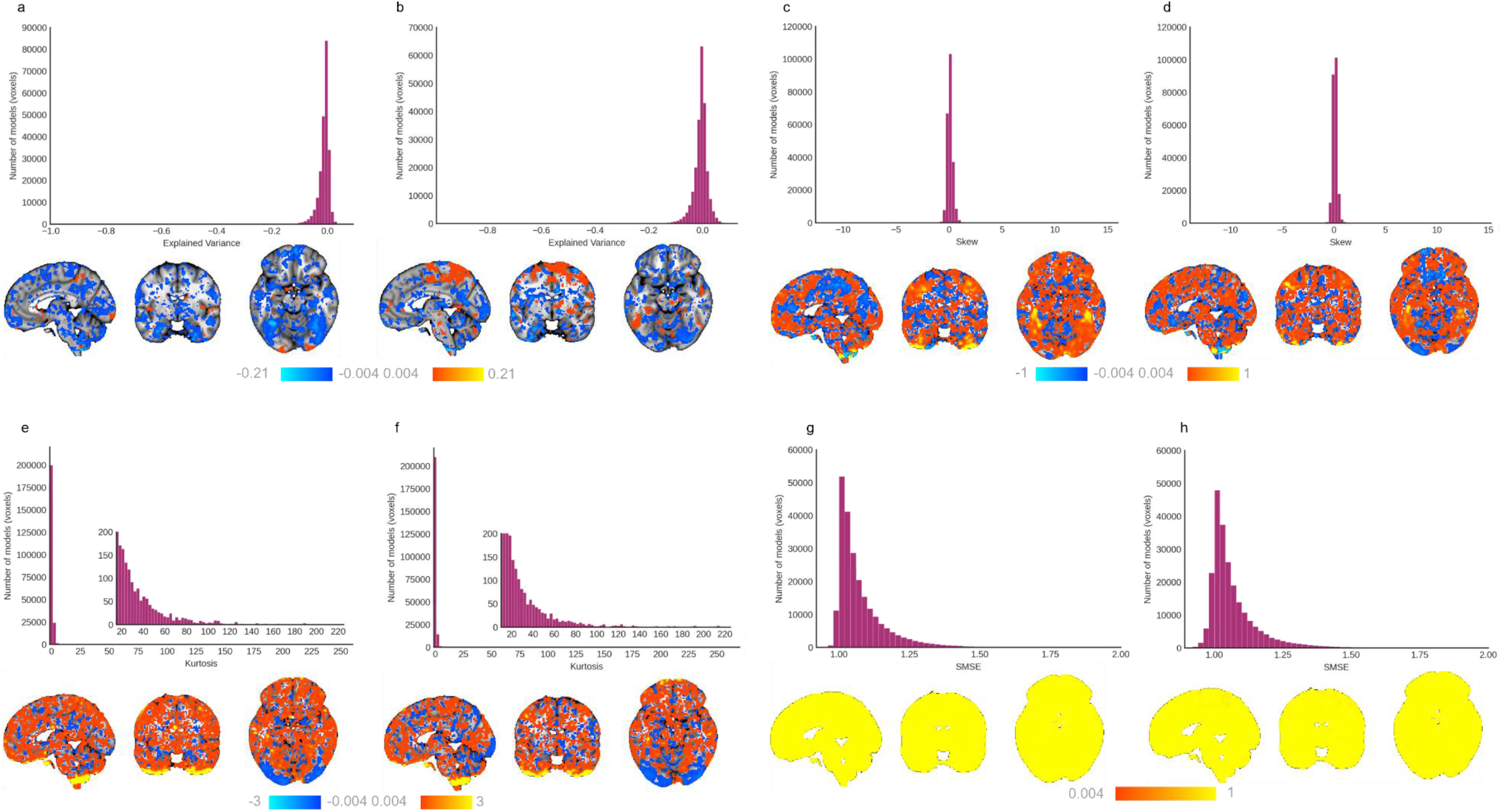
Evaluation of the faces>shapes (left) and faces>baseline normative models when applied to MIND-Set cohort. Histograms show the explained variance (a,b), skew (c,d), kurtosis (e,f), and SMSE (g,h) of the clinical data, as tested on reference normative models of EFMT related BOLD activation, and their respective illustration on the brain. x,y,z= 4,-6,-16.

### SUPPLEMENTARY RESULTS – *Location of deviations for diagnoses*

**Supplementary Figure 8:**
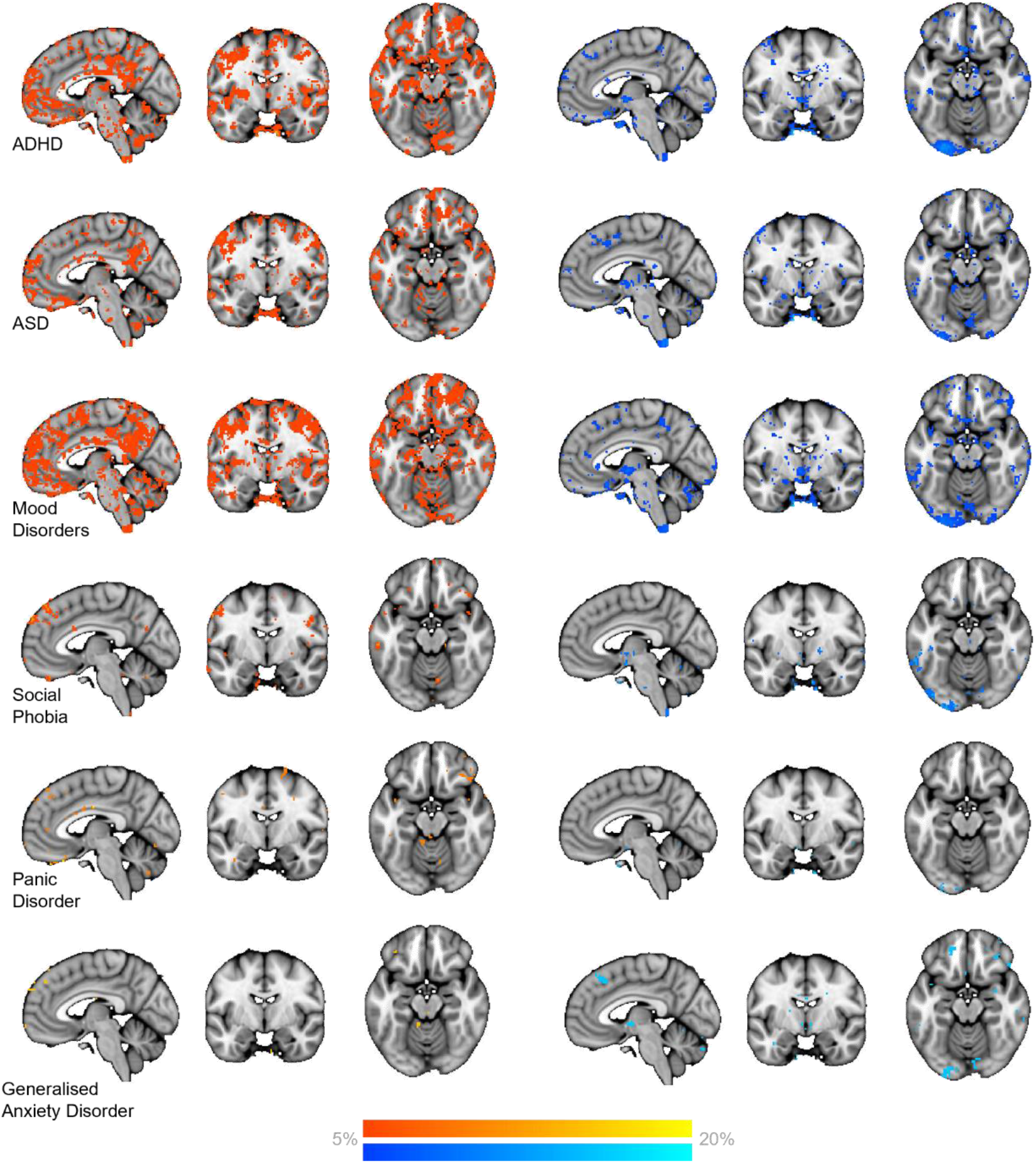
Heterogeneous location of deviations in predicted BOLD signal for different types of neurodivergence, and mental health diagnoses. Maps illustrate the percentage of participants with a neurodivergence or mental health condition who had positive (left; hot colours) or negative deviations (right; cool colours) > ±2.6 within each voxel [minimum = %5 of sample, or 1 participant where 5% was a participant count less than 1, maximum = 20% of disorder sample size]. x,y,z, = -4,- 6,-16.

### SUPPLEMENTARY RESULTS – *Location of deviations for increasing levels of co-occurring diagnoses*

**Supplementary Figure 9:**
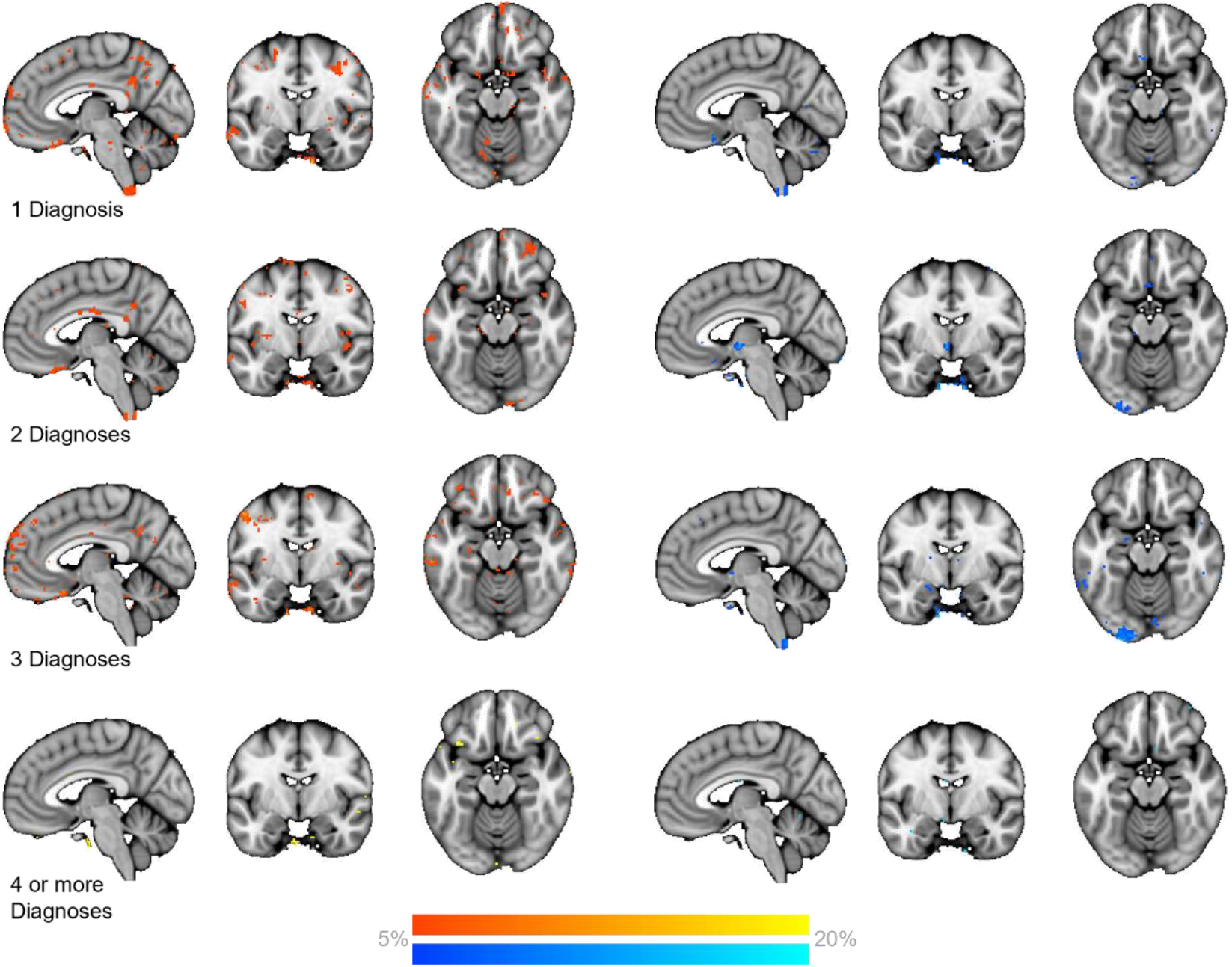
Heterogeneous location of deviations in predicted BOLD signal for increasing levels of co-occurring diagnoses. Maps illustrate the percentage of participants with a neurodivergence or mental health condition who had positive (left; hot colours) or negative deviations (right; cool colours) > ±2.6 within each voxel [minimum = %5 of sample, or 1 participant where 5% was a participant count less than 1, maximum = 20% of sample size]. x,y,z = -4,-6,-16.

**Supplementary Figure 10:**
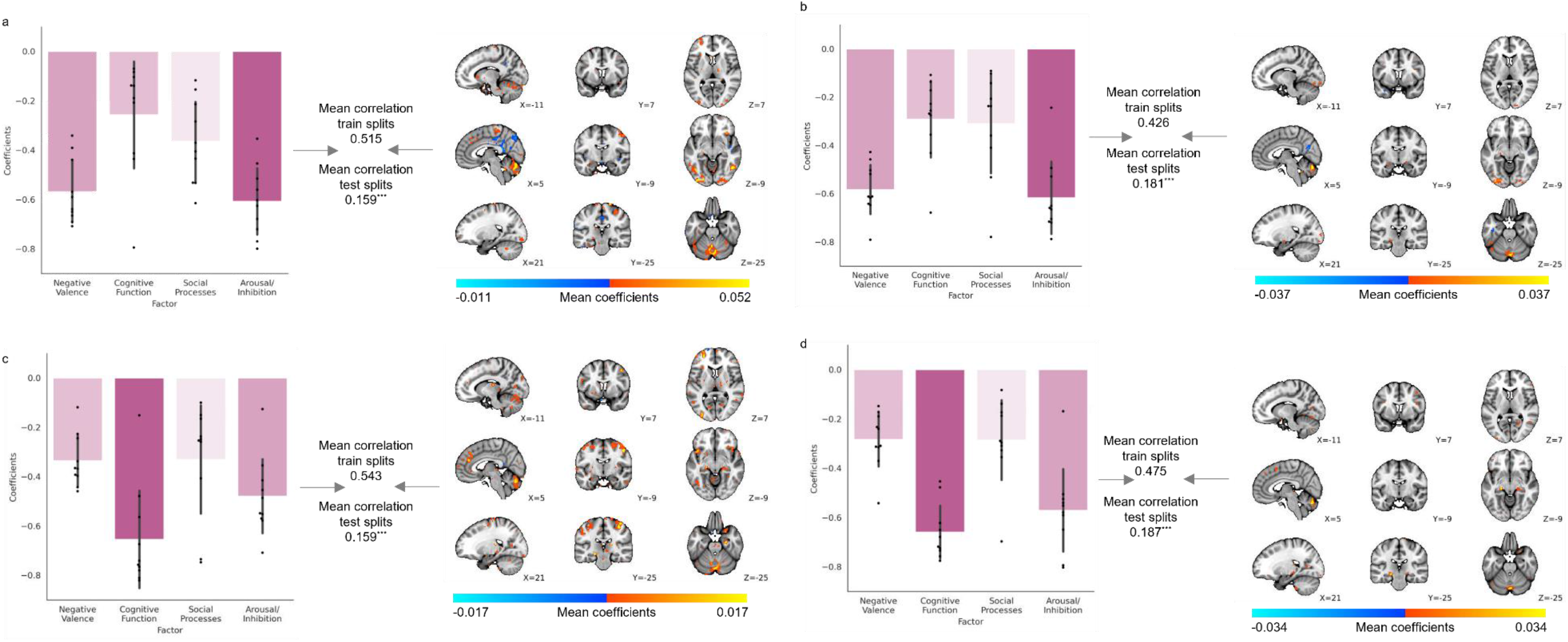
Sparse canonical correlation analyses (SCCA) between functional domains, and deviation scores from faces>shapes or faces>baseline normative models constrained to grey matter (left) and task positive regions (right). Weights per factor to latent variable of psycho-social functioning, canonical correlation between 4 functional domains and whole-brain deviation scores, and their relative mean voxel-wise weights to latent variable of deviation scores from masked (a,b) faces>shapes, and (c,d) faces>baseline normative models (regularisation 10%). All results are statistically significant with 1000-fold permutation tests (*** = *p*<0.001).

## References

1 Marquand, A. F. et al. Conceptualizing mental disorders as deviations from normative functioning. Molecular Psychiatry 24, 1415–1424, doi:10.1038/s41380-019-0441-1 (2019).

2 Marquand, A. F., Rezek, I., Buitelaar, J. & Beckmann, C. F. Understanding Heterogeneity in Clinical Cohorts Using Normative Models: Beyond Case-Control Studies. Biological psychiatry 80, 552–561, doi:10.1016/j.biopsych.2015.12.023 (2016).

3 Rutherford, S. et al. Charting brain growth and aging at high spatial precision. eLife 11, e72904, doi:10.7554/eLife.72904 (2022).

4 Zabihi, M. et al. Fractionating autism based on neuroanatomical normative modeling. Translational Psychiatry 10, 384, doi:10.1038/s41398-020-01057-0 (2020).

5 Zabihi, M. et al. Dissecting the Heterogeneous Cortical Anatomy of Autism Spectrum Disorder Using Normative Models. Biological psychiatry. Cognitive neuroscience and neuroimaging 4, 567–578, doi:10.1016/j.bpsc.2018.11.013 (2019).

6 Bethlehem, R. A. I. et al. A normative modelling approach reveals age-atypical cortical thickness in a subgroup of males with autism spectrum disorder. Communications Biology 3, 486, doi:10.1038/s42003-020-01212-9 (2020).

7 Wolfers, T. et al. Mapping the Heterogeneous Phenotype of Schizophrenia and Bipolar Disorder Using Normative Models. JAMA Psychiatry 75, 1146–1155, doi:10.1001/jamapsychiatry.2018.2467 (2018).

8 Cropley, V. L. et al. Brain-Predicted Age Associates With Psychopathology Dimensions in Youths. Biological Psychiatry: Cognitive Neuroscience and Neuroimaging 6, 410–419, 10.1016/j.bpsc.2020.07.014 (2021).

9 Hariri, A. R., Tessitore, A., Mattay, V. S., Fera, F. & Weinberger, D. R. The amygdala response to emotional stimuli: a comparison of faces and scenes. Neuroimage 17, 317–323, doi:10.1006/nimg.2002.1179 (2002).

10 Hariri, A. R. et al. Serotonin Transporter Genetic Variation and the Response of the Human Amygdala. 297, 400–403, doi:doi:10.1126/science.1071829 (2002).

11 Miller, K. L. et al. Multimodal population brain imaging in the UK Biobank prospective epidemiological study. Nat Neurosci 19, 1523–1536, doi:10.1038/nn.4393 (2016).

12 Van Essen, D. C. et al. The WU-Minn Human Connectome Project: an overview. Neuroimage 80, 62–79, doi:10.1016/j.neuroimage.2013.05.041 (2013).

13 Van Essen, D. C. et al. The Human Connectome Project: a data acquisition perspective. Neuroimage 62, 2222–2231, doi:10.1016/j.neuroimage.2012.02.018 (2012).

14 Harms, M. P. et al. Extending the Human Connectome Project across ages: Imaging protocols for the Lifespan Development and Aging projects. Neuroimage 183, 972–984, doi:10.1016/j.neuroimage.2018.09.060 (2018).

15. Snoek, L., et al. The Amsterdam Open MRI Collection, a set of multimodal MRI datasets for individual difference analyses. Scientific Data 8, 85, doi:10.1038/s41597-021-00870-6 (2021).

16 van Eijndhoven, P. et al. Measuring Integrated Novel Dimensions in Neurodevelopmental and Stress-Related Mental Disorders (MIND-SET): Protocol for a Cross-sectional Comorbidity Study From a Research Domain Criteria Perspective. JMIRx Med 3, e31269, doi:10.2196/31269 (2022).

17 Mulders, P. C. R. et al. Striatal connectopic maps link to functional domains across psychiatric disorders. Translational Psychiatry 12, 513, doi:10.1038/s41398-022-02273-6 (2022).

18 Witten, D. M., Tibshirani, R. & Hastie, T. A penalized matrix decomposition, with applications to sparse principal components and canonical correlation analysis. Biostatistics 10, 515–534, doi:10.1093/biostatistics/kxp008 (2009).

19 Witten, D. M. & Tibshirani, R. J. Extensions of Sparse Canonical Correlation Analysis with Applications to Genomic Data. 8, doi:doi:10.2202/1544-6115.1470 (2009).

20 Bayer, J. M. M. et al. Accommodating site variation in neuroimaging data using normative and hierarchical Bayesian models. NeuroImage 264, 119699, 10.1016/j.neuroimage.2022.119699 (2022).

21 Nygaard, V., Rødland, E. A. & Hovig, E. Methods that remove batch effects while retaining group differences may lead to exaggerated confidence in downstream analyses. Biostatistics 17, 29–39, doi:10.1093/biostatistics/kxv027 (2016).

22 Kebets, V. et al. Fronto-limbic neural variability as a transdiagnostic correlate of emotion dysregulation. Translational Psychiatry 11, 545, doi:10.1038/s41398-021-01666-3 (2021).

23 Rutherford, S. et al. Evidence for embracing normative modeling. eLife 12, e85082, doi:10.7554/eLife.85082 (2023).

24 Westlin, C. et al. Improving the study of brain-behavior relationships by revisiting basic assumptions. Trends in cognitive sciences, doi:10.1016/j.tics.2022.12.015 (2023).

25 Everaerd, D., Klumpers, F., Oude Voshaar, R., Fernández, G. & Tendolkar, I. Acute Stress Enhances Emotional Face Processing in the Aging Brain. Biological Psychiatry: Cognitive Neuroscience and Neuroimaging 2, 591–598, 10.1016/j.bpsc.2017.05.001 (2017).

26 Mizzi, S., Pedersen, M., Lorenzetti, V., Heinrichs, M. & Labuschagne, I. Resting-state neuroimaging in social anxiety disorder: a systematic review. Molecular Psychiatry, doi:10.1038/s41380-021-01154-6 (2021).

27 Elliott, M. L. et al. What Is the Test-Retest Reliability of Common Task-Functional MRI Measures? New Empirical Evidence and a Meta-Analysis. Psychological Science 31, 792–806, doi:10.1177/0956797620916786 (2020).

28 Somerville, L. H. et al. The Lifespan Human Connectome Project in Development: A large-scale study of brain connectivity development in 5-21 year olds. Neuroimage 183, 456–468, doi:10.1016/j.neuroimage.2018.08.050 (2018).

29 Alfaro-Almagro, F. et al. Image processing and Quality Control for the first 10,000 brain imaging datasets from UK Biobank. Neuroimage 166, 400–424 (2018).

30 van Eijndhoven, P. F. P. et al. Measuring Integrated Novel Dimensions in Neurodevelopmental and Stress-related Mental Disorders (MIND-Set): a cross-sectional comorbidity study from an RDoC perspective. medRxiv, 2021.2006.2005.21256695, doi:10.1101/2021.06.05.21256695 (2021).

31 Oldehinkel, M. et al. Attention-Deficit/Hyperactivity Disorder symptoms coincide with altered striatal connectivity. Biological psychiatry. Cognitive neuroscience and neuroimaging 1, 353–363, doi:10.1016/j.bpsc.2016.03.008 (2016).

32 Glasser, M. F. et al. The minimal preprocessing pipelines for the Human Connectome Project. Neuroimage 80, 105–124, doi:10.1016/j.neuroimage.2013.04.127 (2013).

33 Pruim, R. H. R. et al. ICA-AROMA: A robust ICA-based strategy for removing motion artifacts from fMRI data. Neuroimage 112, 267–277, doi:10.1016/j.neuroimage.2015.02.064 (2015).

34. Botvinik-Nezer, R., et al. Variability in the analysis of a single neuroimaging dataset by many teams. Nature 582, 84-88, doi:10.1038/s41586-020-2314-9 (2020).

35. Andersson, J. L., Jenkinson, M., Smith, S. & Oxford, F. A. G. o. t. U. o. Non-linear registration, aka Spatial normalisation FMRIB technical report TR07JA2. 2, e21 (2007).

36 Kraha, A., Turner, H., Nimon, K., Zientek, L. R. & Henson, R. K. Tools to support interpreting multiple regression in the face of multicollinearity. Frontiers in psychology 3, 44, doi:10.3389/fpsyg.2012.00044 (2012).

